# Single-cell multiomics reveals a gene regulatory circuit promoting leukemia cell differentiation

**DOI:** 10.1101/2024.02.29.582681

**Authors:** Xin Tian, Liuqingqing Zhang, Yijia Tang, Guiqiyang Xiang, Zixuan Wang, Ping Zhu, Shuting Yu, Fangying Jiang, Shuai Wang, Jinzeng Wang, Yao Dai, Desheng Zheng, Jianbiao Wang, Han Liu, Xiangqin Weng, Shengyue Wang, Yun Tan, Zhu Chen, Saijuan Chen, Feng Liu

**Affiliations:** Shanghai Institute of Hematology, State Key Laboratory of Medical Genomics, National Research Center for Translational Medicine (Shanghai), Ruijin Hospital Affiliated to Shanghai Jiao Tong University School of Medicine, Shanghai 200020, China; School of Life Sciences and Biotechnology, Shanghai Jiao Tong University, Shanghai 200240, China; School of Computer Science, Southwest Petroleum University, Chengdu 610500, China; Clinical Laboratory, Ruijin Hospital Affiliated to Shanghai Jiao Tong University School of Medicine, Shanghai 200020, China

**Author notes:** Correspondence: Saijuan Chen; Feng Liu. These authors contributed equally to this work (Xin Tian, Liuqingqing Zhang, Yijia Tang).

**Keywords:** acute myeloid leukemia, all-trans-retinoid acid, differentiation trajectory, gene regulatory circuit, scATAC-seq, single-cell multiomics

## Abstract

Cancer differentiation therapy aims to induce maturation of neoplastic cells, yet how cell fate is determined in oncogenic cellular contexts is poorly understood. Here we integrate chromatin accessibility and transcriptome analysis at single-cell levels to investigate the mechanism driving the differentiation trajectory of the PML/RARα^+^ acute promyeloid leukemia (APL) cell line model, NB4, under the induction of all-trans-retinoid acid (ATRA). We show that differentiating NB4 cells navigate through an intermediate cell fate decision point that leads to either terminal granulopoiesis or a transient immature-lymphocyte-like state. Functional perturbation studies indicate that ATRA signaling activates discrete PML/RARα-target enhancers to induce a positive feedforward gene regulatory circuit involving two myeloid-lineage transcription factors, SPI1 and CEBPE, which is both necessary and sufficient to activate a myelocytic gene expression program in NB4. Further, ectopic expression of SPI1 and CEBPE also promotes myelocytic differentiation in non-APL leukemia cell lines HL60 and K562. Together, these results shed light on the differentiation trajectory of therapy-induced cell decisions of APL and suggest a gene regulatory circuit that may be broadly exploited to promote terminal differentiation of leukemia.

## Introduction

Leukemia-initiating cells (LICs) are neoplastic hematopoietic stem/progenitor cells (HSPCs) with infinite self-renewal capacitates^1^. Inducing LICs to differentiate into mature blood cells has long been recognized as a way to clear leukemia blasts from circulatory system^2^. Recently, a series of studies further showed that converting LICs to specialized myeloid antigen-presentation cells (such as dendritic cells and macrophages) can lead to long-lasting and specific anti-tumor immune responses, possible by enhancing neoantigen presentation and recognition by cytotoxic T lymphocytes^3, 4^. Thus, directed differentiation or cellular reprogramming of leukemia blasts holds the potential to not only abolish leukemogenesis but also serve as the basis for developing novel anti-cancer vaccines^5^.

Because leukemic blasts resemble developmentally arrested HSPCs^2, 6, 7^, normal hematopoietic lineage-determining TFs have been exploited to direct maturation of leukemia blasts with the expectation that leukemic blasts would assume the trajectory of normal hematopoietic cells towards maturation^8^. Indeed, this is the basis for the development of TF cocktails to, for example, reprogram various leukemia cells to nonleukemic macrophages using C/EBP_α_ and PU.1/SPI1^3, 4^ or to classical Dendritic cells using PU.1, IRF8, and BATF3^9^. However, normal hematopoiesis is controlled by densely interconnected transcriptional circuits and the functionality of TFs is critically dependent on the accessible chromatin landscape within the cell. Yet it is often unclear if normal hematopoietic TFs or combinations retain the ability to direct cell differentiation in leukemic cells, whose genomic and epigenomic background notably differ from that of normal cells. As a result, time-consuming trial-and-error screening strategies are still the prevalent approach to define the cell reprogramming factor for the induction of leukemia cell differentiation.

Among WHO-classified AML subtypes, acute promyeloid leukemia (APL) with t(15;17)(q22;q12) chromosomal translocation represents a unique case that can be cured by differentiation therapy^10, 11^. The leukemic state of APL cells is transformed and maintained by PML/RAR_α_, an oncogenic fusion transcription factor, while all-trans retinoid acid (ATRA) and arsenic trioxide (ATO) can modulate the gene regulatory function and/or stability of PML/RAR_α_, leading to terminal differentiation of APL blasts towards mature granulocyte-like cells^10^. Given the robust differentiation phenotype of APL *in vitro* and *in vivo* ^12, 13^, we reasoned that its underlying differentiation trajectory could serve as the basis for identifying key differentiation drivers in a leukemic cell context. To that end, we previously performed scRNA-seq analysis to characterize how NB4, an APL patient-derived t(15;17)^+^ cell line, responded to the induction of ATRA for up to 6 days *in vitro*, and we identified a comprehensive set of TFs that were induced by ATRA over the course of NB4’s differentiation towards granulocytes^14^. Nevertheless, scRNA-seq analysis is essentially rooted in correlational analysis and thus insufficient to definitively link these TFs to ATRA signaling or to the directionality and process of differentiation. Therefore, since then, we have generated chromatin accessibility profiles of ATRA-treated NB4 cells using single-cell Assay for Transposase-Accessible Chromatin using sequencing (scATAC-seq). Here we present our integrated scRNA-seq and scATAC-seq of ATRA-induced differentiation trajectory of NB4, which led to the discovery of a cell fate bifurcation point in this process and a feedforward gene regulatory circuit therein to promote terminal granulocytic differentiation. The importance of this regulatory circuit was tested by a suite of functional perturbation studies in NB4 and other non-APL cells lines (K562 and HL60). Our study demonstrates the usefulness of single-cell trajectory analysis in rationally identifying transcription factors networks that may be used to reprogram the fate of leukemia cells.

## Results

### Integrated single-cell chromatin accessibility and transcriptome analysis of ATRA-induced NB4 cells

A series of scATAC-seq libraries were generated from NB4 cells incubated with 1 μM of ATRA for 0/1/3/6 days using a droplet-based combinatorial indexing protocol^15^. As expected, the scATAC-seq reads were correlated strongly with bulk ATAC-seq data and enriched in cis-regulatory elements (CREs) (**Fig. S1, S2A-2B**). After removing low quality cells and putative doublets (**Fig. S2C**; also see **Methods** for details), a total of 21,088 cells were retained for subsequent analysis (Control, 6412 cells; ATRA_day1, 4006 cells; ATRA_day3, 5385 cells; ATRA_day6, 5285 cells; **Table S1**).

To annotate the scATAC-seq data, we pooled scATAC-seq reads from the ATRA-treatment time course for an unsupervised cell clustering analysis. Projecting the data to lower dimensions using the Uniform Manifold Approximation and Projection (UMAP) method revealed inter- and intra-sample heterogeneities (**Fig. 1A**). For example, the ATRA_day0 sample were distributed in two clusters that notably differed in size, echoing earlier reports that naive NB4 cells were predominantly in an undifferentiated state (C1) while a small fraction (<0.1%) showed spontaneous, though aberrant, differentiation characteristics (C2) (**Fig. 1B**)^14^. Cells after ATRA-induction formed several large clusters clearly separated from untreated cells, with cells from the day1 or day6 samples each forming closely positioned cluster (C3 for day1; C6 and C7 for day6), whereas those from the day3 sample were partitioned into two separated clusters (C4 and C5), suggesting that distinct cell fates were induced during ATRA treatment (**Fig. 1B**).

**Figure 1.**
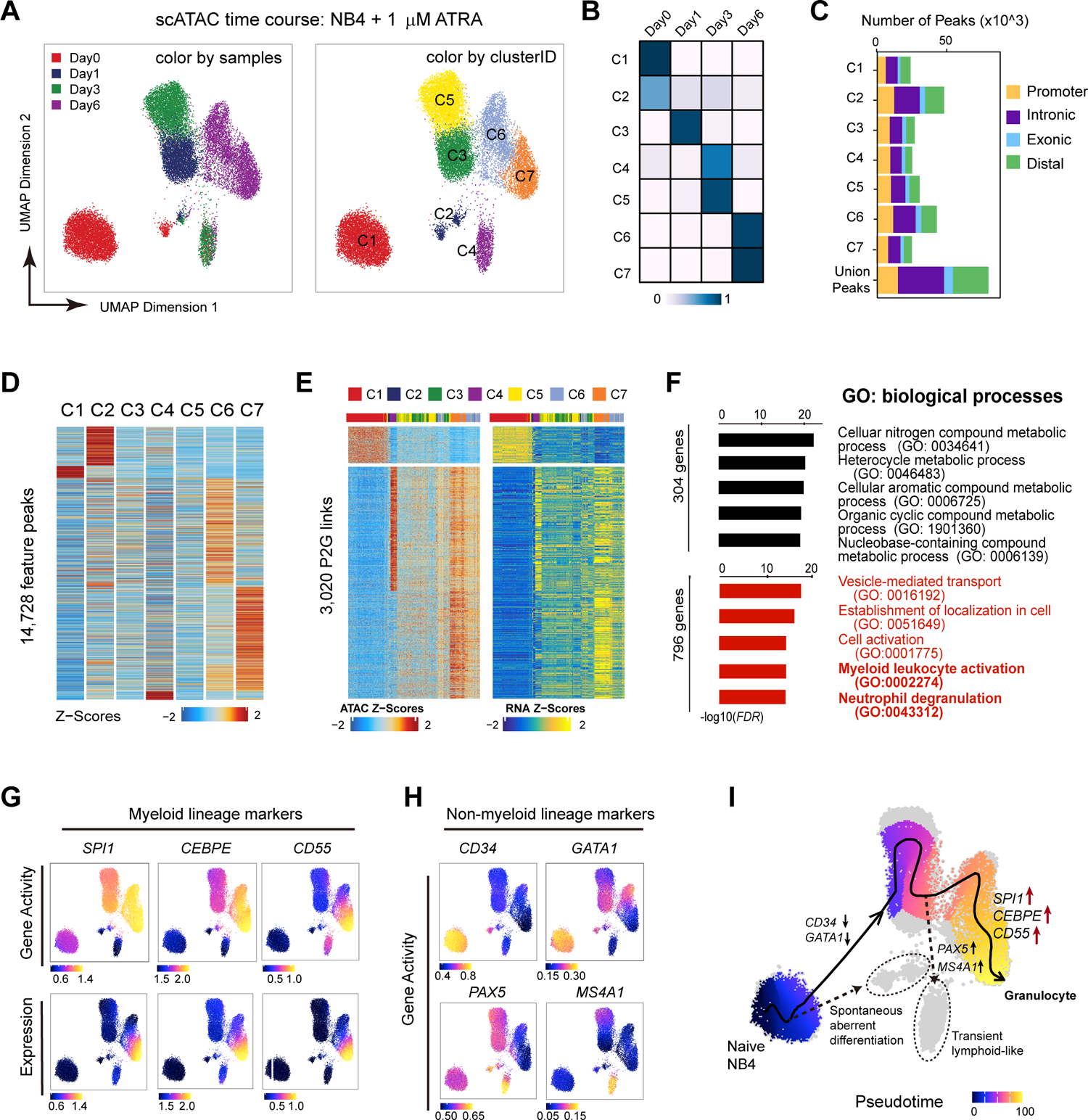
Single-cell multi-omics analysis of the time course of ATRA-induced differentiation A) Uniform manifold approximation and projection (UMAP) plot of scATAC-seq profiles of NB4 cells treated with 1 μM of ATRA for 0/1/3/6 days. Dots represent individual cells; colors indicate sample collection time. Cells were colored by the time of sample collection (left panel) or cluster identity (right panel). B) Heatmap of the fraction of each cluster per experiment. C) Bar chart of peak numbers in each cluster. Colors indicate genomic annotations of peaks. D) Heatmap of feature peaks (cluster-specific peaks). E) Heatmaps of 3,020 peak-to-gene (P2G) linkages identified from integrated matrix of scATAC-seq and scRNA-seq data. F) Gene ontology analysis of down- and up-regulated genes from P2G linkages shown in E. Bar charts of -log_10_(FDR) of top enriched GO-biological processes terms are shown. G) UMAPs of selected myeloid-lineage markers. Color intensities were according to z-scores of gene activities (top row) and gene expressions (bottom row). H) UMAPs of selected non-myeloid-lineage markers. Only the z-scores of gene activities are shown; the expression levels of these genes were not detectable levels in scRNA-seq data. I) UMAP of the pseudotime values of the main trajectory leading to mature granulocyte-like cells. Two side branches were marked in gray: one represents spontaneous and aberrant expression from naïve NB4 cells; the other represents a route digressed the main trajectory, leading to transient lymphoid-like cell state. The dynamic profiles of various lineage-marker genes are illustrated.

Next, we called accessible chromatin peaks on a cluster-by-cluster basis^16^, leading to a union set of 79,658 scATAC-seq peaks (**Fig. 1C; Table S2**). Differential peak analysis yielded 14,728 feature peaks whose accessibilities showed certain cluster-specificities. The accessibility profiles of these peaks appeared to be relatively high in cells collected from day0 (C1-C2) or day6 (C6-C7), but low in cells taken from day1 (C3) or day3 (C4-C5) (**Fig. 1D**), suggesting that the process of ATRA-induction was accompanied by reprogramming of the accessible chromatin landscape.

Last, we applied the Cicero algorithm to compute gene activity (GA) scores genome-wide by aggregating linked distal regulatory elements (enhancers) and promoters at each gene locus^17^. In parallel, we integrated the scATAC-seq data with our earlier scRNA-seq data of the same time course (ATRA treatment for 0/1/3/6 days)^14^ using a semi-supervised inter-dataset alignment approach whereby cells taken from each time point were specified^16^. Unsupervised analysis of the integrated matrix identified 3020 significantly correlated peak-to-gene (P2G) links. These P2G links included 304 down-regulated genes that were mainly associated with cellular metabolic processes, and 796 up-regulated genes associated with myeloid maturation such as myeloid leukocyte activation and neutrophil degranulation (**Fig. 1E-1F; Table S3**). Thus, the integrated single-cell chromatin and transcriptome analysis was able to identify temporally coordinated changes of gene expression programs.

### A cell fate bifurcation point during ATRA-induced NB4 differentiation

It is noteworthy that our earlier clustering analysis using scRNA-seq data alone did not show clear separation of cell fates in the day3 sample—that is, the transcriptomes of cells at day3 were largely grouped together in scRNA-seq-based clustering^14^, which contrasted to the appearance of two distinct clusters in scATAC-based clustering (**Fig. 1A-1B**). Upon comparison of the profiles of GA scores (computed from scATAC-seq data) and gene expressions (computed from scRNA-seq data), we noticed that, at day3, SPI1 and CEBPE’s chromatin activities were already high in C3 but still low in C4 (**Fig. 1G, upper panels**); by contrast, both genes’ expression was equally low in C3 and C4 (**Fig. 1G, lower panels**). Thus, a higher sensitivity of chromatin activities of granulopoiesis genes (as compared to their transcription) in distinguishing intermediate cellular state could underpin the resolution of intra-sample heterogeneity of day3 samples.

In addition, GA scores revealed cluster-specific activities of a panel of well-established non-myeloid marker genes—including those of HSPC (CD34), erythrocyte (GATA1), and B lymphocyte (PAX5 and MS4A1)—even though that the expression levels of these genes were all very low in bulk and single-cell RNA-seq data throughout the course of ATRA-induction (**Fig. 1H; Fig. S3**). More interestingly, the GA scores of non-myeloid marker genes exhibited cluster- and temporal-specificities different from those of myeloid marker genes. For example, the GA scores of CD34 and GATA1 were relatively high in untreated NB4 cells (C1) but were absent from all ATRA-treated cell clusters (C3-C7), suggesting a rapid loss of the chromatin signatures of HSPC or erythrocyte marker genes upon ATRA-induction. The GA scores of PAX5 and MS4A1 were initially low in untreated NB4 cells (i.e., C1) and then elevated specifically in C4, a cluster from the day3 sample. The PAX5^+^/MS4A1^+^ GA signature, however, was absent from cells of day6 sample (i.e., C6 and C7). Together, these results revealed an intermediate cell fate bifurcation point along the differentiation trajectory of ATRA-induced NB4 cells: at about day3 post induction, one groups along the axis of C1-C3-C5-C6-C7 that led to granulocytic differentiation, a group of cells (C4) digressed, towards a transient lymphoid-like state (**Fig. 1I**)

### *De novo* identification of candidate TF drivers of ATRA-induced differentiation

Based on the differentiation trajectories outlined above, we went on to use the integrated scATAC-seq and scRNA-seq data to identify TFs driving the trajectories of 1) C1-C3-C5-C6-C7 (Trajectory-1) that leads to terminal granulocytes and 2) C1-C3-C4 (Trajectory-2) that leads to a transient lymphoid-like state. Given the shared origin of these two fates, we reasoned that TF drivers shared by both trajectories might be involved in an early phase of differentiation (C1-C3), whereas trajectory-specific TF drivers were involved in later phases (C3-C5-C6-C7 or C3-C4) (**Fig. 2A**).

**Figure 2.**
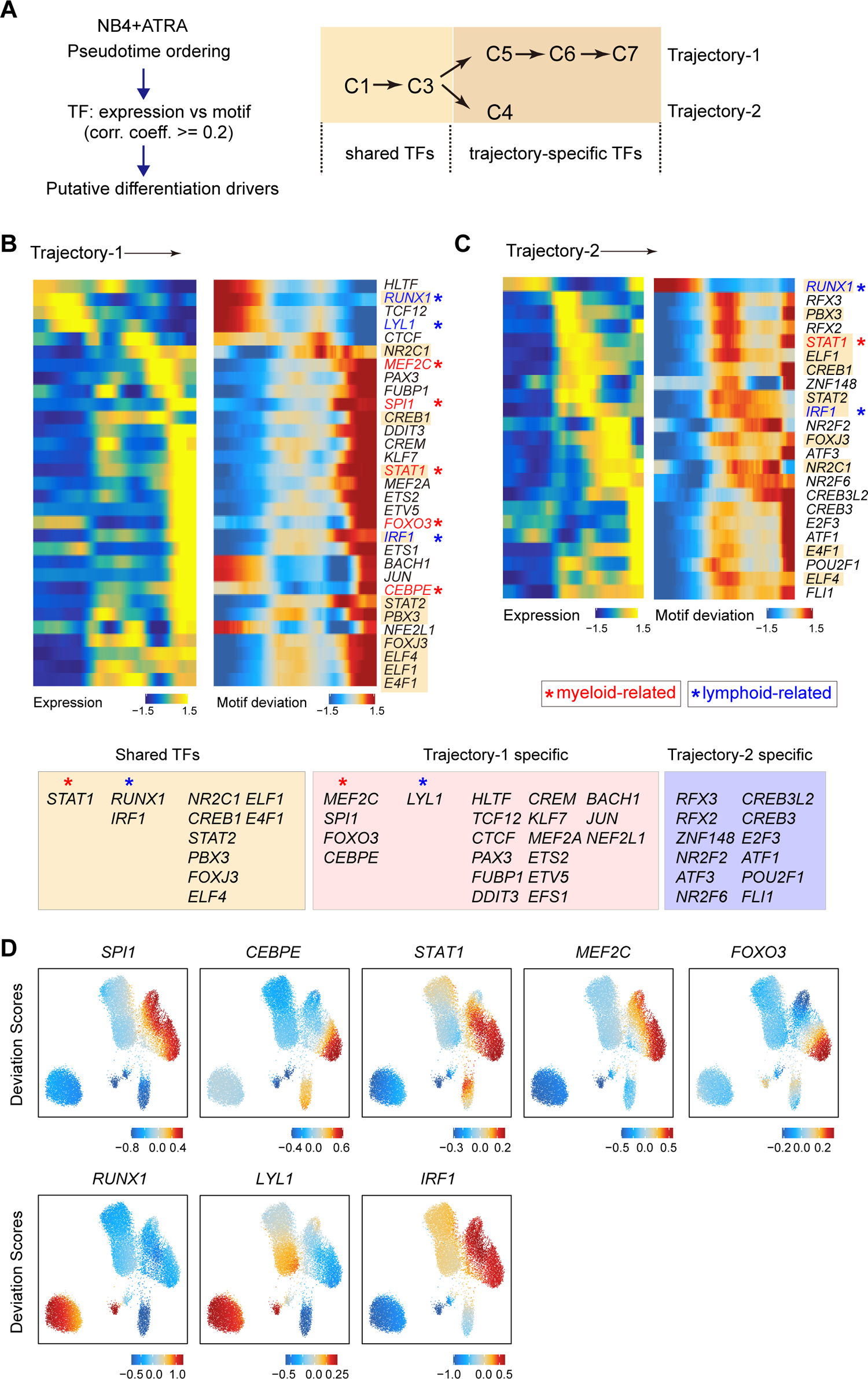
*De novo* identification of the TF drivers of ATRA-induced granulopoiesis A) Schematics of the analysis pipeline to identify differentiation driver TFs. Cells along two trajectories were separately subjected to pseudotime analyses. B-C) Heatmap of pseudotime-dependent peaks and genes in Trajectory 1 (B) and Trajectory 2 (C). Putative driver TFs shared in both trajectories are shaded. Red and blue indicate factors related to myeloid- or lymphoid-development GO terms, respectively. D) UMAPs of selected TFs. Colors indicate the TF’s motif deviation scores.

For the identification of candidate TF drivers, we used a strategy proposed by Granja et. al.,^16^ to compute the correlation co-efficiencies between the expression of TFs and their motif deviation scores. Setting the cutoff of expression-*vs*-motif correlation coefficiency at 0.2 (pairwise *Pearson correlation* test, *p* < 0.05) led to 31 and 23 candidate TF drivers for Trajectory-1 and −2, respectively (**Fig. 2B-2C**). Of them, 11 TFs were shared between trajectories, including three (RUNX1, STAT1, and IRF1) that were associated with myeloid (GO:0061515) or lymphoid (GO:0030098) development. Traejectory-1-specific driver TFs included four myeloid-related TFs (i.e., SPI1, CEBPE, MEF2C, and FOXO3; all up-regulated) and one lymphoid-related TF (LYL1, down-regulated), whereas none of the Traejectory-2-specific TF drivers were known to be associated with myeloid or lymphoid development (**Fig. 2B-2C**). In support of a role for Trajectory-1-specific TFs in driving the progression of ATRA-induced granulocyte-differentiation, the motif deviation scores for several Trajectory-1 specific TFs—including SPI1 and CEBPE—showed gradual and cluster-specific increases along the trajectory towards granulocytes (**Fig. 2D**).

### Dynamic reprogramming of PML/RAR**α**-target enhancer networks during ATRA-induced differentiation

ATRA-signaling is known to act through the transcription factor function of PML/RAR_α_ and PML/RAR_α_-binding elements were proposed as ATRA-responsive enhancers to induced the expression of differentiating-related genes^18^. To characterize the dynamics of enhancers along the trajectory of ATRA-induced differentiation, we applied the Cicero algorithm^17^ on the scATAC-seq data, identifying 4,671,794 co-accessible peak pairs, with an average of 1,458,959 positively co-accessible peak pairs per cluster (range: 1,335,060-1,663,462). These co-accessible peaks were further classified to cis-co-accessibility networks (CCANs), which were chromosome-wide modules of highly co-accessible sites indicative of coordinated transcriptional activities during cell differentiation^17^ (**Fig. 3A**).

**Figure 3.**
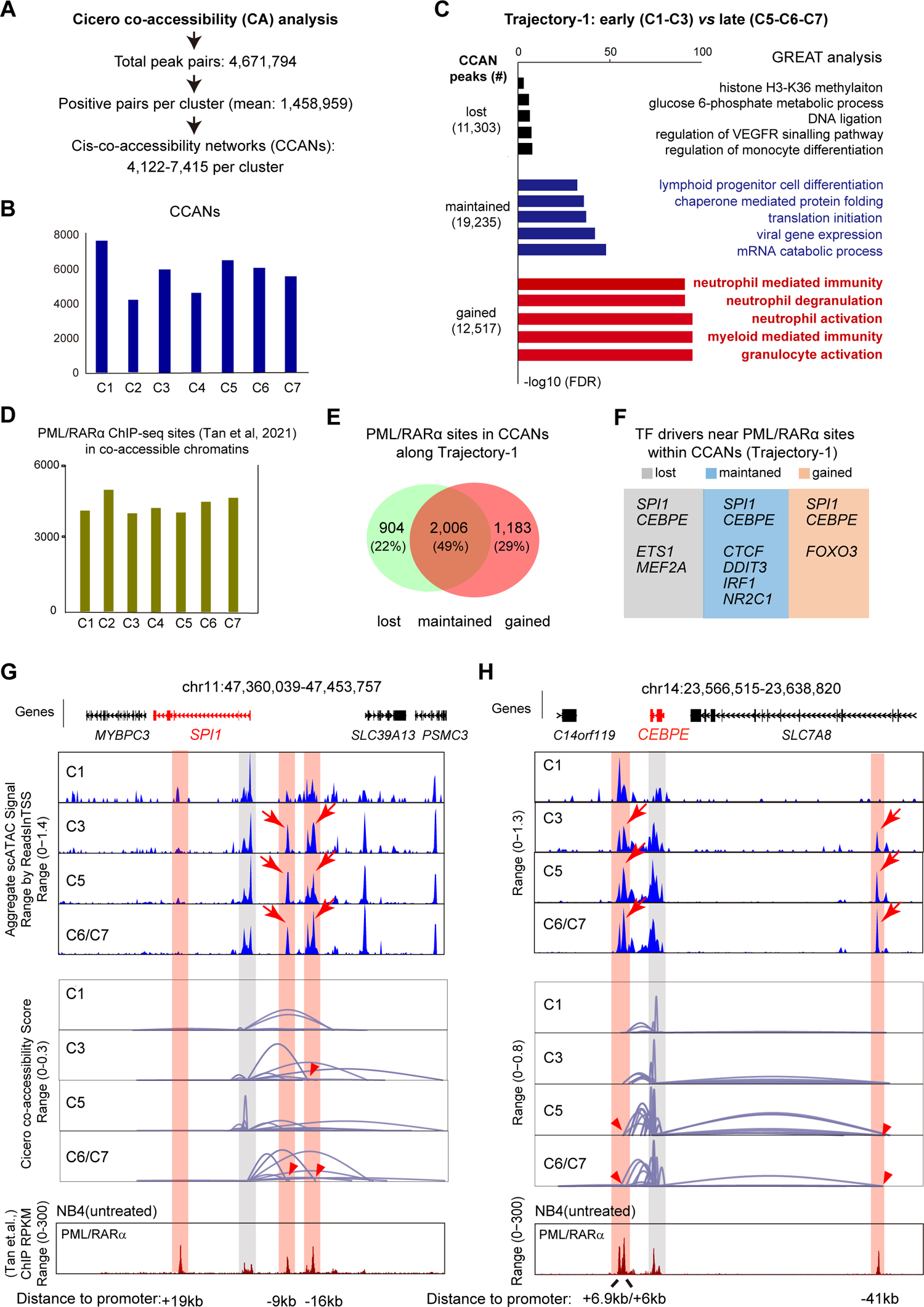
Cis-co-accessibility-network analysis reveals genome-wide reprogramming of enhancer networks A) Flowchart of Cicero analysis to identify co-accessible (CA) peaks and cis-co-accessibility networks (CCANs). B) Bar chart of CCANs per cluster. C) Dynamics of accessible peaks during early-to-late phase transitions in Trajectory-1. The peaks that were lost, maintained, or gained during phase transitions were used as input for GREAT analysis. Bar charts of -log10 (FDR) are shown. GO-biological process terms are at right. D) Bar chart of the distribution of PML/RARα ChIP-seq sites overlapping with CA peaks per cluster. E) Venn diagram of PML/RARα ChIP-seq sites in dynamic CCAN peaks during phase transitions in Trajectory-1. F) Putative TF drivers nearest to dynamic PML/RARα ChIP-seq sites during phase transitions in Trajectory-1. G) Track plots of aggregated scATAC-seq peaks (blue tracks at top panels), cis-co-accessibility links (grey tracks in the middle panels) and PML/RARα ChIP-seq peaks (red tracks at the bottom panels) at the SPI1 (A) and CEBPE (B) loci. The positions of SPI1 or CEBPE promoters are shaded in grey. Adjacent strong PML/RARα-binding sites are shaded in light red. The sgRNA targets for CRISPRi experiments are indicated below. Base-pair distances are with respect to SPI1 or CEBPE transcription start sites (TSS). kb: kilobases.

We then examined the membership of accessible peaks included in CCANs in different phases of a given differentiation trajectory^17^, with an focus on 6,636 high quality PML/RARα-binding sties determined by published ChIP-seq experiments^19^. Along ATRA-induced granulopoiesis trajectory (Trajectory-1), 11,303, 19,235, and 12,517 peaks were respectively lost, maintained, or gained in CCANs during the transition between an early phase (C1-C3) and a late phase (C5-C6-C7). According to GREAT analysis^20^, the gained CCAN accessible chromatins in this trajectory were significantly associated with granulocyte activation, neutrophil degranulation, and neutrophil mediated immunity, consistent with the expectation that they coordinately regulated granulocytic differentiation (**Fig. 3B-3C**). Moreover, the overall number of PML/RARα-binding sites within co-accessible peaks were similar among clusters (**Fig. 3D**). Yet, of 4,093 PML/RARα-binding sites identified in accessible chromatins along granulopoietic Trajectory-1, 22%, 49% and 29% sites were lost, maintained, and gained in CCANs, respectively, suggesting a dynamic reprogramming of PML/RARα-target enhancers in phase transitions (**Fig. 3E**). By contrast, when a similar analysis for Trajectory-2 leading to the side branch (early phase: C1-C3; late phase: C4), the gained CCAN accessible chromatins towards the side branch were not associated with granulocyte maturation, and of 3,813 PML/RARα-binding sites found along this trajectory, a much larger fraction (52%) of PML/RARα-binding sites were lost from CCANs during phase transition (**Fig. S4**). These results thus suggested that trajectory-specific gene regulatory activities involved reprogramming of distinct cis-regulatory elements.

Furthermore, some dynamic PML/RARα-binding sites along Trajectory-1 were found to be located near several candidate TF drivers (**Fig. 3F**), of which two TFs (SPI1 and CEBPE) appear especially regulated since they were near PML/RARα-binding sites within lost, maintained, and gained CCANs during NB4 differentiation (**Fig. 3F**). A close view at these two gene loci showed multiple PML/RARα-binding sites whose co-accessibility profiles with nearby promoters correlated with ATRA-induced differentiation. That is, two PML/RARα-target sites near SPI1 (−9 kb and −16 kb) and three near CEBPE (−41 kb, +6 kb and +6.9 kb) increasingly became co-accessible with SPI1 or CEBPE’s promoter post ATRA-induction (**Fig. 3G-3H; Fig. S5**). Of them, only the −41 kb PML/RARα site upstream of CEBPE was apparently within an apparently “closed” chromatin before ATRA-induction, whereas the others were already “open” in treatment-naive cells (i.e., those in C1). Thus, ATRA signaling led to both activation of newly accessible enhancers and reprogramming of already accessible enhancers.

### Loss-of-functional mutagenesis of SPI1 or CEBPE interrupts ATRA-induced differentiation

The above TF driver analysis and CCAN analyses, taken together, also suggested a model whereby ATRA signaling directly induced SPI1 and CEBPE via distinct PML/RARα-target enhancers to drive terminal granulocytic differentiation of NB4 cells. To functionally interrogate this model, we first performed CRISPR-Cas9-mediated loss-of-functional (LOF) assays to evaluate the role of both TFs in ATRA-induced differentiation.

Specifically, we generated LOF mutant clones carrying frame-shift nonsense mutations in the coding sequences of either SPI1 or CEBPE (**Fig. 4A**). These clones were treated with 1 μM of ATRA for 0/1/3/6 days followed by RNA-seq. As shown in the principal component analysis (PCA) plot, the transcriptomes of wild type (WT) NB4 cells during ATRA-induction were aligned along a curve that was indicative of their differentiation trajectory (**Fig. 4B, left panel**). The transcriptomes of SPI1 or CEBPE mutants induced by ATRA for 0/1/3 days all clustered closely with their counterpart control experiments, whereas the mutants induced for 6 days appeared to “lagged” behind their counterpart WT cells along the differentiation axis (**Fig. 4B, middle and right panels**). Consistent with this, Euclidean distances between transcriptomes revealed a gradual decrease of similarity between ATRA-induced WT NB4 cells and SPI1/CEBPE mutants (**Fig. 4C**). Differentially expressed gene (DEG) analysis indicated that many ATRA-induced genes failed to be up-regulated in SPI1 or CEBPE mutants between day3 and day6 (**Fig. 4D, Table S6**), concomitant with a loss of myeloid leukocyte differentiation (GO:0002573) gene signature at day6 (**Fig. 4E**). Wright-Giemsa stain further showed that most ATRA-induced WT NB4 cells contained condense multi-lobed nuclei characteristic of mature granulocytes, but SPI1 or CEBPE mutants largely retained immature-like nuclei (large and round) even after 6 days of ATRA-induction (**Fig. 4F**). These results provided molecular and phenotypic evidence that SPI1 or CEBPE were responsible for driving ATRA-induced granulopoiesis.

**Figure 4.**
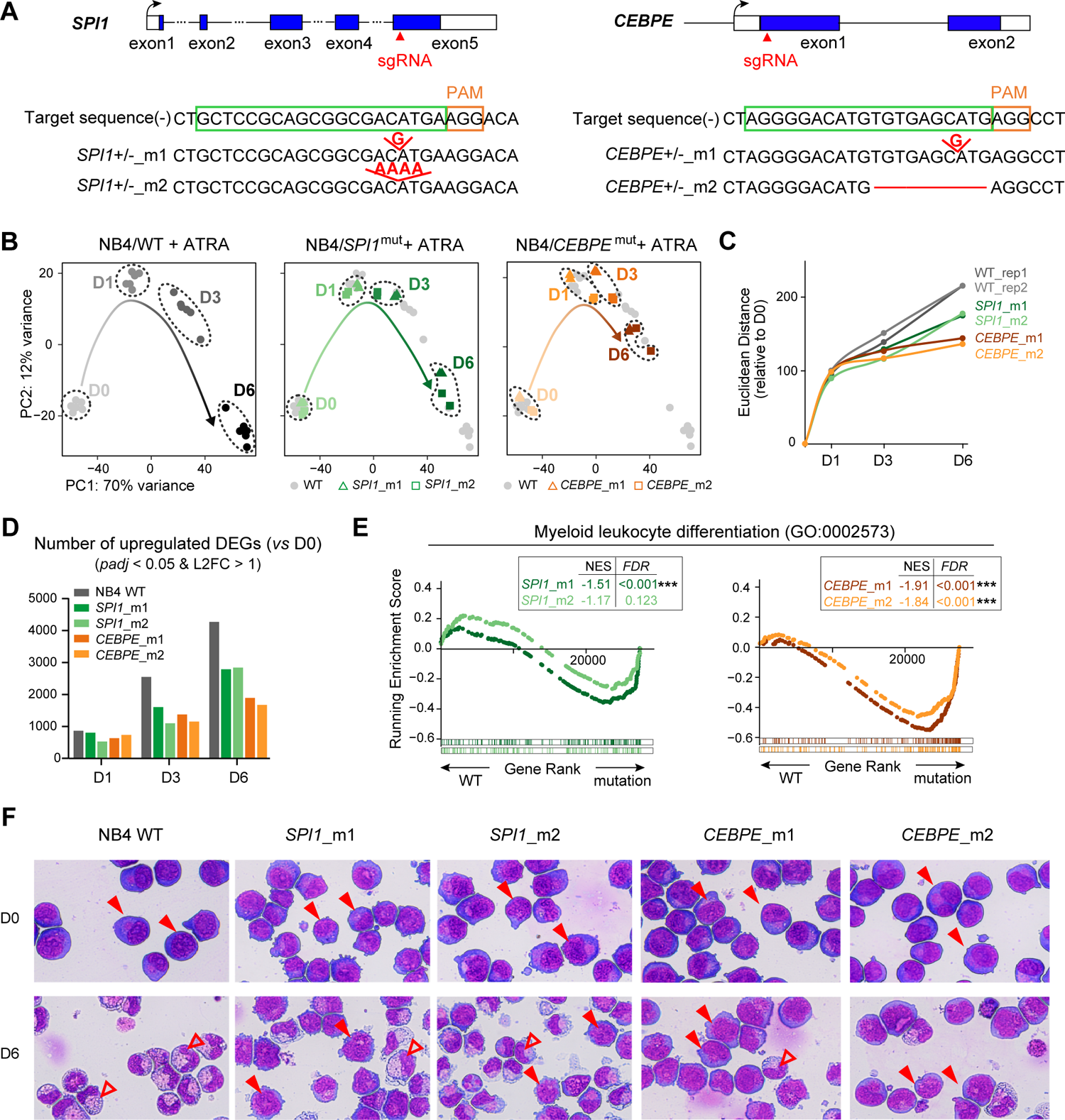
Mutations in SPI1 and CEBPE interrupt ATRA-induced differentiation in NB4 cells. A) Schematics of CRISPR/Cas9-mediatd mutagenesis of SPI1 and CEBPE. Positions of sgRNAs were shown. For each gene, two mutant clones were generated. Deleted or inserted nucleotides are indicated in red. B) PCA plots of unsupervised transcriptome analyses. Each dot represents an RNA-seq experiment. The left panel shows control experiments using WT NB4 cells. The middle and right panels respectively show experiments of mutants of SPI1 (green) and CEBPE (orange), while the control experiments are shown in the background in light grey. C) Euclidean distances between ATRA-induced transcriptomes and un-induced transcriptomes shown in (B). D) Numbers of upregulated differentially expressed genes (DEGs) between ATRA-induced cells and un-induced (D0) cells. E) GSEA analysis of myeloid-leukocyte-differentiation-related genes in SPI1- and CEBPE-mutant cells after 6 days of ATRA-induction. F) Wright-Giemsa stains of WT or mutant NB4 cells. Filled arrowheads indicate nuclei with blast-like characteristics. Empty arrowheads indicated condensed, multi-lobed nuclei characteristic of mature granulocytes.

### Suppressing PML/RAR**α**-target enhancers interrupts ATRA-induced differentiation

To verify the regulatory linkages between ATRA signaling and SPI1 and CEBPE, we used CRISPR interference (CRISPRi) technology to recruit the dCas9-KRAB fusion protein, an artificial transcription repressor^21^, to six dynamic PML/RARα-binding sites near SPI1 or CEBPE (the relative distance of CRISPRi targets with respective to promoters are shown in **Fig. 3G-3H, bottom row**). We established stable CRISPRi cell lines and treated them with 1 μM of ATRA for 0/1//3/6 days. RNA-seq analysis showed that SPI1 expression was notably reduced in two CRISPRi cells (sg-SPI1^sg1/-^ ^16kb^ and sg-SPI1^sg2/-9kb^) at day3 and day6 post ATRA-induction, indicating these target sites as strong enhancers; in comparison, a minor loss of SPI1 expression was observed for the third cell line (sg-SPI1^sg3/+19kb^), suggesting it as a weak enhancer (**Fig. 5A, left panel**). In a similar vein, the +6 kb PML/RARα-binding site was identified as a strong enhancer for CEBPE, whereas two other sites (−41 kb and +6.9 kb) showed little enhancer activity (**Fig. 5B, left panel**). The observed effects on SPI1’ and CEBPE’s expression are highly specific, because genes near targeted sites showed little changes in these CRISPRi experiments (**Fig. S6A-6B**).

**Figure 5.**
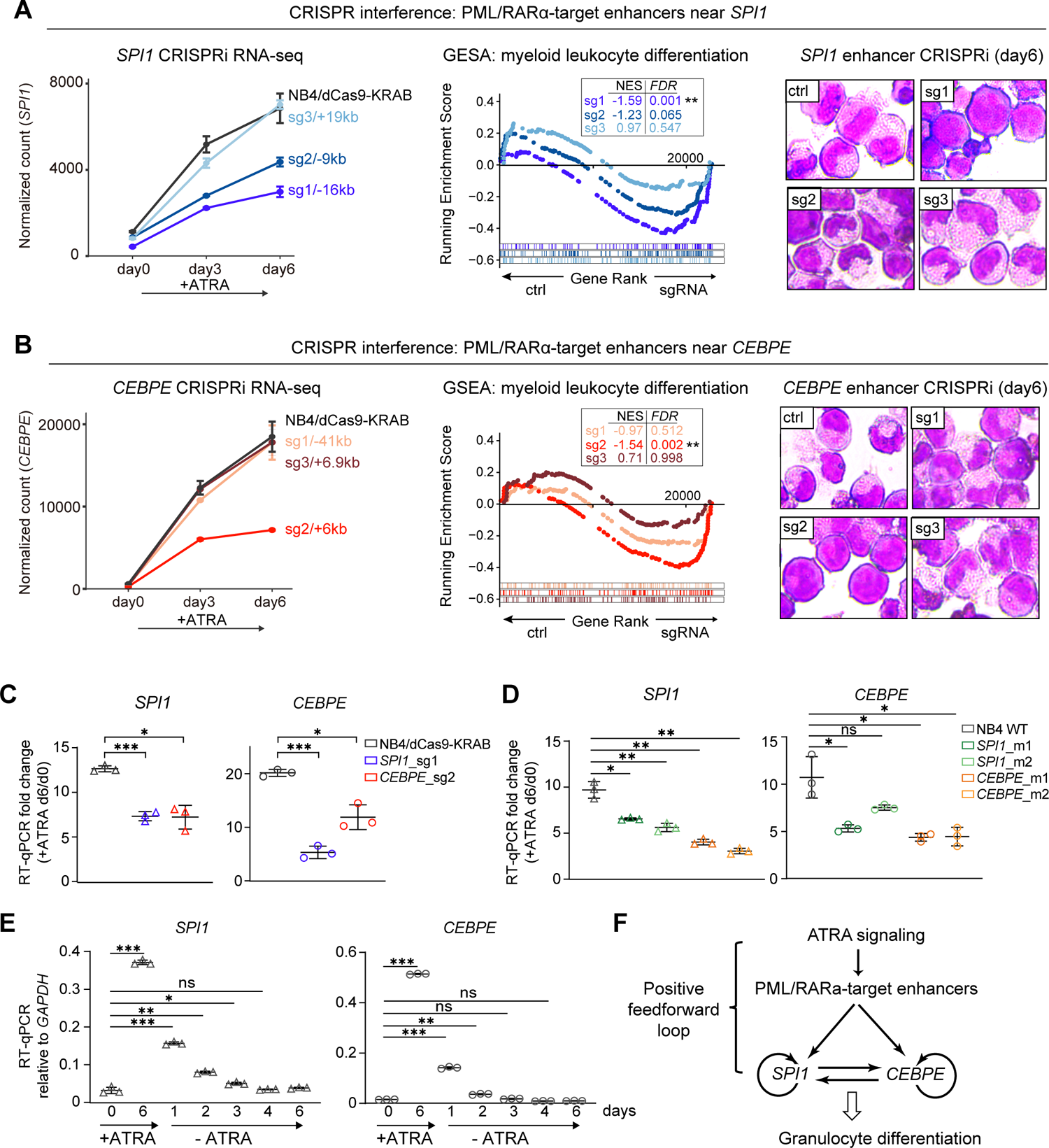
ATRA activates a positive feedforward gene regulatory circuit to drive NB4 differentiation A-B) Line plots of normalized RNA-seq counts of SPI1or CEBPE (left panel), GSEA analysis (middle panel), and Wright-Giemsa stains (right panel) of ctrl(NB4/dCas9-KRAB) or CRISPRi cell lines targeting PML/RARα-sites near SPI1 (A) or CEBPE (B). C) RT-qPCR of indicated CRISPRi cell lines. Internal reference gene is GAPDH. D) RT-qPCR of indicated CRISPR-mediated mutagenesis cell lines. Internal reference gene is GAPDH E) RT-qPCR of NB4 cells in ATRA withdrawal experiments; these cells were first induced with ATRA for 6 days and then cultured in ATRA-free medium for indicated days. F) Schematics of the positive feedforward circuit between ATRA signaling and the expression of SPI1 and CEBPE. Arrows indicate positive regulation. *t*-test: *: *p* < 0.05; **: *p* < 0.01; ***: *p* < 0.0001.

Like what was observed in SPI1 or CEBPE LOF mutants, transcriptome similarity, GSEA, and Wright-Giemsa staining all showed that ATRA-induced NB4 differentiation were interrupted, albeit to various degrees, in some CRISPRi cell lines (**Fig. 5A-5B, middle and right panels; Fig. S6C-6D, S7**). The change of transcriptome profiles was most notable when strong enhancers were targeted (i.e., sg-SPI1^sg1/-16kb^, and sg-CEBPE^sg2/+6kb^), and less so when weaker enhancers were targeted (SPI1^sg2/-9kb^ and SPI1^sg3/+19kb^). DEGs not upregulated between CRISPRi cells and their control counterparts (NB4/dCas9-KRAB) largely overlapped and were significantly associated with myeloid differentiation and activation (**Fig. S6E**). These results indicated that ATRA signaling acted through specific PML/RARα-target enhancers to elevate the expression of SPI1 and CEBPE, which in turn promoted NB4 cells to differentiate towards granulocytes.

### Cross-regulation between SPI1 and CEBPE establishes an interlinked regulatory circuit

Curiously, although SPI1 and CEBPE are respectively located on the 11^th^ and 14^th^ chromosomes and enhancers usually act in cis^22^, knocking down either SPI1’s or CEBPE’s enhancers led to transcriptional loss of both genes post ATRA-induction, a phenomenon that was confirmed by quantitative reverse transcription PCR (RT-qPCR) (**Fig. 5C**). RT-qPCR also showed that both SPI1 and CEBPE’s expressions were significantly reduced in NB4 cells carrying LOF mutations of each gene (**Fig. 5D**). And when NB4 cells were first incubated with 1 μM of ATRA for six days and then in ATRA-free medium for additional 1-6 days, both SPI1 and CEBPE’s transcription levels gradually fell to levels similar to those of untreated NB4 cells (**Fig. 5E**).

These results are indicative of a two-layered feedforward gene regulatory circuit, whereby ATRA signaling mobilizes distinct PML/RARα-target enhancers to induce the expression of SPI1 and CEBPE; and when both TFs’ levels rise above certain threshold levels, they self- and cross-regulate each other to further escalade each other’s levels to activate a terminal granulopoiesis program (**Fig. 5F**). Such intercalated gene regulatory circuit fulfilled the definition of a type of positive feed-forward gene network motif that is frequently utilized for cell fate specification and/or differentiation^23^.

### The SPI-CEBPE regulatory circuit mediates cell fate transitions of NB4 cells

Recently, McKenzie et al. showed that fate of ATRA-induced APL blasts is plastic and could reverse back to immature states upon ATRA-withdrawal^24^. To determine if such fate plasticity is mediated by the SPI1-CEBPE gene regulatory circuit, we analyzed the transcriptomes of NB4 WT and CRISPRi cells pre-treated by ATRA for 1, 2, 4, or 6 days and followed by culture in ATRA-free medium for another 3 days (**Fig. 6A**). In the transcriptome-based PCA plot, ATRA-withdrawal cells—including those pre-induced for 1/2/4/6 days—seemingly formed a distinct cluster separated from those during ATRA-induction. In comparison, when strong ATRA-responsive SPI1 and CEBPE’s enhancers were knocked down (i.e., CRISPRi:SPI1_sg1 and CRISPRi:CEBPE_sg2), the resultant transcriptomes were grouped more closely with untreated NB4 cells (**Fig. 6B**). DEG analysis indicated 1,368 genes that were relatively insensitive to ATRA-withdrawal in WT NB4 cells, approximately half of which fell to near baseline levels post ATRA-withdrawal in CRISPRi:SPI1_sg1 and CRISPRi:CEBPE_sg2 cells (**Fig. 6C**). Thus, the relative insensitivity of some genes to ATRA-withdrawal could be attributed, at least in part, to the regulatory circuit established by ATRA-responsive, PML/RARα-target enhancers.

**Figure 6.**
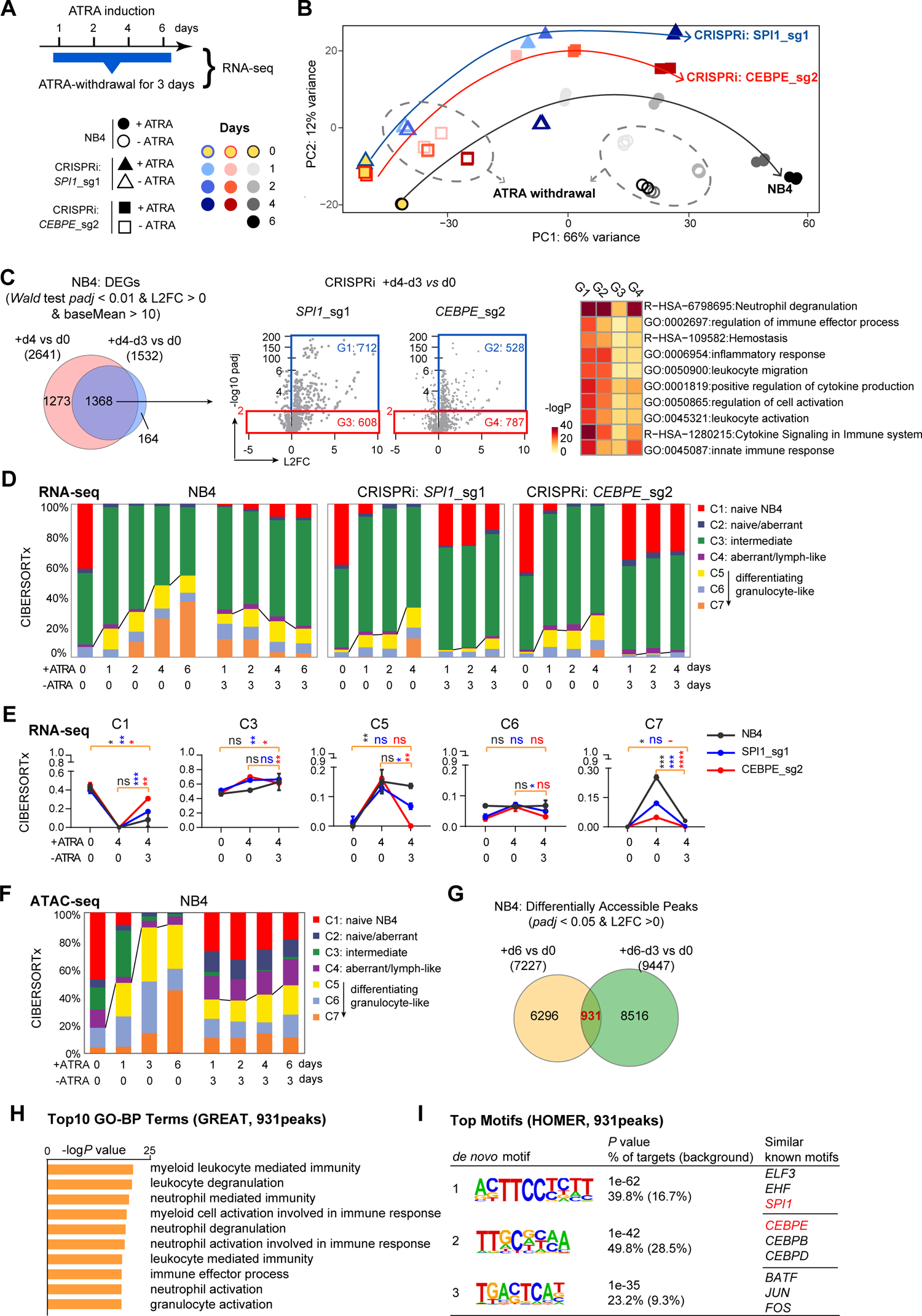
ATRA-responsive PML/RAR**α**-target enhancers are required for cell fate commitment of NB4 cells A) Experimental scheme ATRA withdrawal experiment, whereby the control cell line (dCas9-KRAB) or CRISPRi NB4 cell lines (dCas9-KRAB + enhancer-targeting sgRNAs) were induced by ATRA for various days and then cultured in ATRA-free medium for three days (ATRA-withdrawal). B) PCA plots of unsupervised transcriptome analyses. The legend is shown in (6A). Each dot represents an RNA-seq experiment. Samples corresponding to ATRA withdrawal experiments circled by dashed lines. C) Venn diagram of the result of DESeq2 analysis of ATRA-induced (+d4) or ATRA-withdrawal (+d4-d3) experiments, as compared to untreated cells (d0). 1,368 ATRA-induced DEGs were did not fall back to baseline levels in these experiments (left panel). About half of these genes, however, were reduced to baseline levels in CRISPRi cells targeting strong SPI1- or CEBPE-enhancers (middle panel). GO terms associated with DEGs responding to ATRA-withdrawal were enriched with immune responses and granulocyte maturation (only GO terms with *padj* < 0.01 were shown) D) CIBERSORTx deconvolution of bulk RNA-seq experiments in (A). Colored bars represent relative fractions of the gene expression signatures of single-cell-analysis-derived cell clusters/states. E) Line plots of the relative fractions of each cluster-specific signature in bulk RNA-seq experiments of dCas9-KRAB NB4 and CRISPRi cells. Days of ATRA-induction and ATRA-withdrawal are indicated below. F) CIBERSORTx deconvolution of ATAC-seq experiments. Colored bars represent relative fractions of the signature chromatin accessibility profiles of single-cell-analysis-derived cell clusters/states. G) Venn diagram of differentially accessible peaks in ATAC-seq experiments. ATRA-treated cells for 6 days with or without 3 days of ATRA-withdrawal (+d6-d3 or +d6) are compared to untreated cells (d0). H-I) GREAT and motif analyses of 931 ATRA-induced peaks that did not fall back to baseline level of accessibility in ATRA-withdrawal experiments.

We further applied the CIBERSORTx algorithm^25^ to deconvolute the bulk RNA-seq data of ATRA-withdrawal cells using cluster-specific gene expression profiles derived from integrated scRNA-seq and scATAC-seq data (**Fig. 6D; Fig. S8; Table S4**). As expected, during ATRA-induction, the fractions of the expression signatures of C1 (immature cell state) decreased whereas the signatures of more mature granulocyte-like cell clusters/states (C5/C6/C7) gradually increased over time (**Fig. 6D**). In ATRA-withdrawal transcriptomes, there was a trend of gene-expression-signature-profiles regressing to that of untreated cells (C1), though the signature of C5, largely remained unchanged. However, when in CRISPRi cells (in which SPI1- and CEBPE-enhancers were knocked down), C5’s signature decreased to baseline levels post ATRA-withdrawal (**Fig. 6E**). Because C5 represents an intermediate cluster/state just after the cell fate bifurcation point wherein SPI1 and CEBPE began to upregulated post ATRA-induction (**Fig. 2D**), its dependence on distinct ATRA-responsive enhancers supported the notion that these enhancers were responsible for establishing or maintaining the intermediate cell state towards terminal granulocytic differentiation.

Deconvolution analysis of bulk ATAC-seq data of ATRA-treated time course led to a similar conclusion. The accessible landscape of ATRA-withdrawal cells did not regress to that of untreated cells and were characterized by an enlarged representation of the features of several intermediate clusters (especially C5) (**Fig. 6F**). 931 accessible chromatin peaks were found to be initially activated by ATRA but were insensitive to ATRA-withdrawal and these peaks were strongly associated with myeloid differentiation and activation and most highly enriched with binding motifs for SPI1 and CEBPE (**Fig. 6G-6I**), consistent with the idea that SPI1-CEBPE-regulatory network mediate the commitment of ATRA-induced NB4 cells.

### Transgenic SPI1 and CEBPE expression broadly promotes myelocytic differentiation in leukemia cells

Finally, we assessed the possibility to use the SPI1 and CEBPE-regulatory circuit to promote leukemia differentiation in the absence of ATRA. To that end, we generated doxycycline (Dox)-inducible plasmid constructs for either or both of SPI1 and CEBPE in NB4 and two other PML-RARα^-^ leukemia cell lines: HL60 (an AML-M2 subtype-like cell line) and K562 (an BCR-ABL1^+^ chronic myelogenous leukemia cell line). These cell lines were incubated with Dox for 4 days to induce SPI1 and/or CEBPE expression. To compare the effect of ATRA signaling and TF over-expression (OE), the parental WT cell lines were also independently treated with 1 μM of ATRA. The induced levels of SPI1 and CEBPE were quantified by RT-qPCR and transcriptomes of these experiments were profiled by DRUG-seq, a plate-based high throughput 3’ RNA-seq protocol^26, 27^ (**Fig. 7A-7B**).

**Figure 7.**
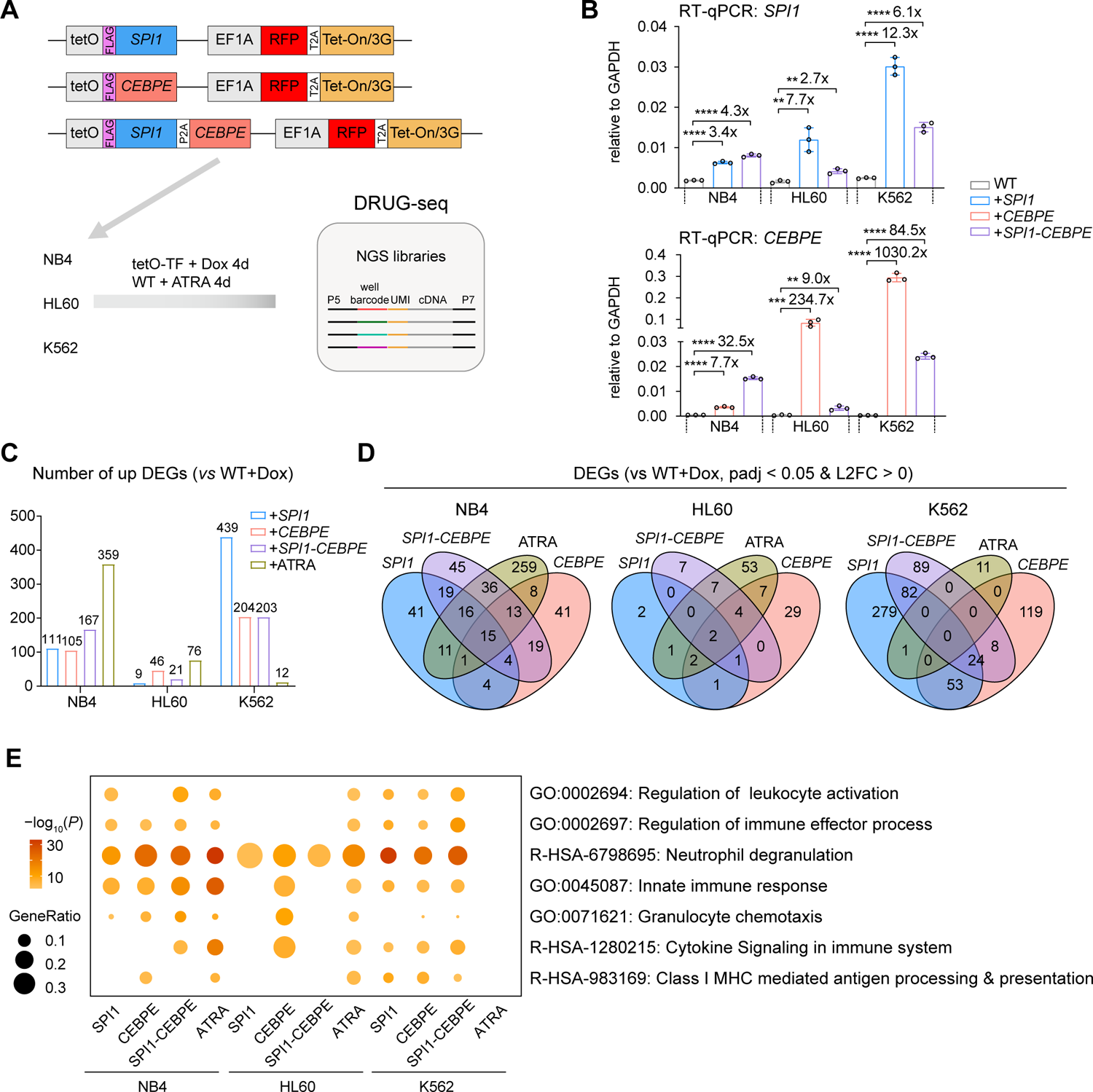
SPI1 and CEBPE expression drives leukemia cell lines to differentiate A) Study design overview. Inducible SPI1/CEBPE or dual TFs expression genes were inserted into genome of leukemia cell lines by lentivirus infection. After induction with doxycycline (Dox) for 4 days, cells were collected and prepared for sequencing following DRUG-seq protocol. B) RT-qPCR for detection of SPI1/CEBPE expression level after doxycycline induction for 4 days. C) Bar plot of upregulated genes in each group compared with their wild type (WT) after doxycycline induction for 4 days. D) Venn diagram of upregulated genes after TF overexpression or ATRA treatment in three cell lines. E) Enrichment of upregulated genes in each group. The color indicates -log_10_(P) value. The size of bubble indicates the number of genes enriched in this term dividing the number of input genes. GO or HSA terms are shown in the right.

In total, we obtained 61 DRUG-seq transcriptomes that passed quality control. Consistent with previous reports, DRUG-seq only generated 3’-end cDNA libraries and thus the detected DEGs were less than those from bulk RNA-seq analysis of NB4 and HL60 cells that we published before^14^ (**Fig. 7C**). Nevertheless, the DRUG-seq data confirmed a cell type-specific effect of ATRA on these cell lines—it induced most genes in NB4 cells while having smaller or little effect on HL60 and K562 cells, respectively (**Fig. 7C**). In NB4, ATRA-induced genes overlapped with 39%, 35%, and 48% of DEGs upregulated in SPI1-, CEBPE-, and dual SPI1/CEBPE-OE cell lines, respectively, and TF-OE-upregulated genes were associated with GO terms related to granulocyte maturation, consistent with the idea that both TFs mediate ATRA-induced granulocytic differentiation in this cell line (**Fig. 7D, Table S7**). In HL60 and K562, TF-OE-upregulated genes showed varying degrees of granulocyte maturation such as neutrophil degranulation. This is especially significant in K562 cells, which was almost completely insensitive to ATRA-induction (**Fig. 7E**).

## Discussion

In this study, we took a bottom-up approach to analyze the gene regulatory network mediating cell fate decisions in NB4, a classical cell line model for differentiation therapy of PML/RARα^+^ APL. Through integrated analysis of two complementary single-cell datasets (scATAC-seq and scRNA-seq) sampled from a time course of ATRA-induced NB4 cells, we uncovered a previously unknown cell fate decision point, after which only a subset of cells with successful upregulation of two differentiation factors, SPI1 and CEBPE, embarked on a trajectory towards terminal differentiation, whereas the rest digressed toward a transient immature lymphoid-like fate. We further showed that SPI1 and CEBPE, as well as their downstream granulopoiesis program, formed an interlinked gene regulatory circuit to commit the cell fate decisions. Last, we demonstrated that, even in the absence of ATRA signaling, transgenic over-expression of SPI1 and CEBPE could activate a myelogenous gene expression program in both APL (NB4) and non-APL (HL60 and K562) leukemia cell lines.

On one hand, the discovery of the feedforward linkages from ATRA signaling to PML/RARα-target enhancers and then SPI1 and CEBPE provided mechanistic insights on the specificity, efficacy, and plasticity of ATRA-induced differentiation of PML/RARα^+^ APL cells. First, because each enhancer is activated by a particular combination of the trans-regulatory inputs^28, 29^, the ATRA-responsive PML/RARα enhancers near SPI1 and CEBPE ensure that both TFs, as well as their downstream gene programs, are up-regulated only when both ATRA and PML/RARα are present. Second, due to dose-dependent actions of both TFs, the differentiation program was only activated by strong ATRA-responsive PML/RARα enhancers. Third, there was a continuous requirement for ATRA in maintaining the enhancers’ activities, so that the differentiation program was partially reversed upon ATRA-withdrawal, providing a mechanistic explanation for the fate plasticity of NB4 cells.

On the other hand, the ability of SPI1 and CEBPE to promote myelocytic differentiation suggests them as potential therapeutic targets to manipulate cell fates in leukemic blasts. This may be especially useful in the context where ATRA is not effective or no exogenous signaling molecules are available as effective differentiation inducers. Proof-of-concept examples were shown here using K562 and HL60, the former an erythrocytic CML cell line that is almost completely ATRA-insensitive (this study) and the latter a FAB-M2-like leukemia cell line in which ATRA only induces incomplete and promiscuous differentiation ^14^. In both cell lines, we observed significant induction of granulocytic gene expressions post SPI1 and/or CEBPE overexpression, suggesting that the SPI1 and CEBPE combination may activate granulocytic gene program in very different leukemic cellular contexts. We note, however, that the cell reprogramming effect is relatively small on HL60. This may suggest the involvement of certain factors negating the function of SPI1 or CEBPE in this cell line, which requires further investigation.

As recent studies have demonstrated, converting leukemic blasts to innate immune cells with elevated antigen presenting capacities might stimulate the production of lymphocytes that can recognize neoantigens, ensuing specific and lasting clearance of leukemia blasts^4^. To screen the TF cocktails for directed differentiation, earlier studies typically started from TFs in control of normal hematopoiesis. However, our study suggests that, in a neoplastic cell context, the TFs with fate-converting capacities might differ from those acting in normal development. For example, in normal development, SPI1/PU.1 is responsible for myeloid commitment beyond myeloblast stage, whereas CEBPE (mouse homolog: C/EBP_ε_) is expressed later to direct last step(s) of neutrophil development such as the formation of secondary or tertiary granule^24, 30, 31^. In contrast, SPI1 and CEBPE appear to be simultaneously up-regulated by ATRA in NB4 cells and then co-regulate a terminal differentiation program, suggesting that they form an equal partnership, rather than a hierarchical relationship, in directing NB4’s differentiation. Therefore, single-cell regulatory trajectory analysis uncovered an otherwise unusual TF combination that was operational in the context of leukemia cell differentiation. Further, the strong PML/RARα-target enhancers mediating the transcriptional induction of SPI1 and CEBPE that we functionally validated may additionally serve as the basis to develop anti-leukemia gene therapies using epigenome-editing therapies^32^.

## Acknowledgement

This study is supported by research funding from National Natural Science Foundation of China (81970130 and 81770143), National Key Research and Development Program of China (2018YFA0107802), Shanghai Commission of Science and Technology (17PJ1405800), Shanghai Municipal Education Commission Gaofeng Clinical Medicine Grant (20171902), and Shanghai Dong Fang Scholarship.

## Conflict of interest

The authors declare that they have no conflict of interest.

## Data availability

Next-generation sequencing data generated in this study are available from publicly accessible database upon publication of this work. Previously published bulk ATAC/RNA-seq and scRNA-seq data of ATRA-treated NB4 cells^14^ are publicly available from The National Omics Data Encyclopedia (https://www.biosino.org/node, accession# OEP001921).

## Author Contributions

FL and SJC conceived and supervised the study. XT, LQQZ, YJT, GQYX, ZXW, PZ, STY, FYJ, SW, YD, JZW, XQW performed experiments and data analysis. DSZ, HL, JBW, SYW, YT analyzed the data. FL, SJC, ZC, XT, LQQZ wrote the paper with comments from all other authors.

## Materials and Methods

### Cell culture

NB4 cells were gifts from M. Lanotte^33^. Cells were cultured in Iscove’s Modified Dulbecco’s Medium (IMDM) supplemented with 10% fetal bovine serum (FBS) (MOREGATE), 100 units/mL penicillin and 100 mg/mL streptomycin (GIBCO) at 37 °C with 5% CO2. ATRA (Brand name: Tretinoin; SelleckChem, Cat# S1653) was dissolved in DMSO as 1 mM stock solution. For ATRA-induction experiments, the ATRA stock was diluted to 1 μM in cell culture medium for indicated times. Mycoplasma tests were routinely performed during this study. For ATRA-withdrawal experiment, the cells were first treated with 1 μM of ATRA for designated days, and then washed with PBS three times before culturing in fresh ATRA-free medium.

### RNA-seq

Total RNA was extracted from 10^6^ to 10^7^ cells with the Qiagen RNAeasy Mini kit (Cat# 74104). RNA libraries were generated from 1-2 μg of total RNA per sample with the VAHTS Stranded mRNA-seq Library Prep Kit following the manufacturer’s recommendations (Vazyme, Cat# NR612). Multiplexed libraries were mixed and sequenced by Illumina NovaSeq6000 platform at the depth of 30-50 million 150 bp paired-end reads per sample. Sequencing reads of each library were pseudoaligned to a transcriptome index file (based on Homo_sapiens.GRCh38.cdna.all.fa) using Kallisto (v0.46.0)^34^. Transcript compatibility counts were used for differential gene expression analyses using the DESeq2 R package (version 4.1.0)^35^. Counts normalized by DESeq2’s estimateSizeFactors and plotCounts function were used to be compared between samples. Counts normalized by varianceStabilizingTransformation function were used for principal component analysis by BiocGenerics (v0.40.0) and Euclidean distances computation by stats R package (v4.1.0). For deconvolution analysis of bulk RNA-seq, matrices of normalized TPM counts and scRNA-seq counts of cells projected to scATAC-seq-defined clusters were uploaded to CIBERSORTx web portal (https://cibersortx.stanford.edu/). To generate the signature matrix, “input data type” was specified as “scRNA-seq”; “Min.expression” was set at 0; all other parameters were as default settings.

### DRUG-seq

DRUG-seq was performed according to Li et.al.^26, 27^, with minor modifications. Each well of 96-well plates was pre-dispensed with 150 μL of lysis buffer and 0.1 μmol DRUG-seq reverse transcription (RT) primers, each containing a 10-nt well-barcode. 50k cells per treatment were then added per well. After shaking for 15 min at 900 rpm, 15 μL of cell lysate per well was transferred into new 96-well PCR plates pre-dispensed with 2.75 μL of RT reaction mix per well. After RT at 42 °C for 90 min, the barcoded cDNAs were pooled for DNA purification, 50 ng of which was used for NGS library preparation (Vazyme Cat# TD501). The resultant libraries were sequenced on the Illumina NovaSeq6000 platform with the addition of a custom Read1 primer^26, 27^. Primers used in DRUG-seq libraries preparation and sequencing are in **Table S5**. The sequencing reads were processed with Drop-seq_tools-2.5.3 (https://github.com/broadinstitute/Drop-seq) (RRID:SCR_018142) to extract well-barcodes corresponding to different experiments and align the coding sequences to human transcriptome (GRCh38 gencode.v25.gene.annotation). This resulted in a digital expression matrix (genes x well-barcodes), which was used as input for the DESeq2 R package (version 4.1.0; RRID:SCR_000154) for differential gene expression analysis.

### ATAC-seq

ATAC experiments were performed according to the fast ATAC-seq protocol^36^. Approximately 50,000 cells were used per experiment. Sequencing libraries were prepared by the TruePrep DNA Library Prep Kit V2 for Illumina (Vazyme, Cat #TD501). The libraries were mixed and sequenced by Illumina NovaSeq6000 at the depth of 40-60 million 150 bp paired-end reads per sample. De-multiplexed reads of each sample were mapped to a human genome index file (hg19) using Bowtie2 (v2.2.9) with default settings. Then duplicate reads generated from PCR were removed by SAMtools (v1.3.1; RRID:SCR_002105). Peaks were called by MACS2 (v2.1.1.20160309) with the following parameters: -f BAMPE -g hg -B -q 0.01. After removing ENCODE blacklist regions^37^, an ATAC-seq reads count table was generated using bedtools’ (v2.25.0) multicov function, which was used as input for DEseq2 to determine differentially accessible peaks between samples. Peakset-associated gene ontology enrichment analysis was performed by GREAT (v4.0.4). Motif enrichment analysis was performed by HOMER (v4.11).

### scATAC-seq

scATAC-seq libraries were performed with the Bio-Rad’s SureCell ATAC-Seq Library Prep Kit according to the manufacturer’s instructions (Bio-Rad, Cat# 17004620). Library concentration and fragment size were checked by Agilent 2100 Bioanalyzer. The libraries were sequenced on Illumina NovaSeq6000 platform with an additional ddSEQ Sequencing Primer as read1 primer. Fastq files of the replicates of scATAC-seq experiments were combined and processed by the Bio-Rad ATAC-Seq Analysis Toolkit (https://hub.docker.com/u/bioraddbg/) for reads debarcoding, genome alignment (hg19), alignment quality check, and cell filtering with default settings. BAM files generated from filtered cells were converted to fragment files by Sinto (version 0.7.3.1) for subsequent single cell analysis. Cell filtering and annotations are summarized in **Table S1**.

### Dimensionality reduction and clustering

Fragment files of all scATAC-seq experiments (NB4_control, NB4_ATRA_day1, NB4_ATRA_day3, NB4_ATRA_day6) were processed by the ArchR package (version 1.0.1; RRID:SCR_020982)^16^. Arrow files were generated with the following filtering parameters: minTSS=7.5, minFrags=1000. Doublet removal, dimensionality reduction, clustering, and single cell embedings were performed with corresponding ArchR functions using default parameters.

### scATAC-seq and scRNA-seq data integration

Single-cell RNA-seq experiments of NB4 cells treated with 1 μM of ATRA for 0/1/3/6 days were reported earlier^14^ and were downloaded from The National Omics Data Encyclopedia (NODE; https://www.biosino.org/node) under the accession number OEP001921. The Seurat object of scRNA-seq data was imported into ArchR for scATAC-seq and scRNA-seq integration using a constrained integration method: in both scRNA-seq and scATAC-seq data, clusters corresponding to the time of collection (i.e., ATRA_day0, ATRA_day1, ATRA_day3, or ATRA_day6) in both datasets were specified before running the ArchR addGeneIntegrationMatrix function.

### Peak calling and annotation

An iterative, reproducible peak calling strategy was used to identify scATAC-seq peaks by ArchR: first, cells of each annotated scATAC-seq cluster were used to generate pseudo-bulk replicates by the addGroupCoverages function of ArchR; second, peaks of each cluster were identified by the addReproduciblePeakSet function with MACS2 (version 2.2.7.1). Cluster-specific marker peaks were identified by getMarkerFeatures. For the peakAnnoEnrichment function, a cutoff was set at “*FDR* <= 0.1 & Log2FC >= 0.5”. Transcription factor motif deviation scores were computed by chromVAR (version: 1.14.0)^38^ with the “cisbp” motif collection^32^. Peak-to-gene linkages were identified by the addPeak2GeneLinks function. Pairwise differential test between clusters were computed using the getMarkerFeatures function by specifying the clusters for comparison.

### Pseudotime ordering

Cells along a given differentiation trajectory were selected for pseudotime analysis in ArchR with the addTrajectory function. Correlation between GeneIntgrationMatrix and MotifMatrix was computed by correlateTrajectories (corCutOff = 0.2, varCutOff1 = 0.5, varCutOff2 = 0.5).

### Chromatin co-accessibility networks

The scATAC-seq count matrix of cells in each annotated cluster were processed by the Cicero R package (version: 1.3.4.11) to compute genome-wide co-accessibility scores of scATAC-seq peaks. Matrices were binarized prior to computing chromatin co-accessibility scores using run_cicero(cicero_cds, human.hg19.genome, sample_num = 100). Cis-co-accessibility Networks (CCANs) were identified by the generate_ccans function with default settings. Promoter-centered view of co-accessibility links were shown for selected genes. Different Cicero CCAN maps were matched using the maxmatching R package (version: 0.1.0) according to Pliner et.al.^17^.

### CRISPR mutagenesis

CRISPR-mediated mutagenesis was carried out by transiently introducing CRISPR/Cas9 RNA-nucleoprotein complex into cultured NB4 cells with the 4D-Nucleofector system (Lonza). The sample preparation kit was Lonza SF Cell Line 4D-Nucleofector Kit S (Cat# V4XC-2024). Each reaction included 4×10^5^ cells, 20 pmol of Cas9-C-NLS nuclease (GeneScript, Cat# Z03385), and 24 pmol of sgRNA (ordered in GeneScript Corp, China) in 20 μL of volume. 48 hours after electroporation, cells were transferred to a 12-well culture plate for expansion. Single-cell clones were obtained by diluting cells and culturing in 96-well plates. For each CRISPR mutagenesis experiment, targeted DNA fragment in 20-30 clones were checked by Sanger sequencing (Genewiz, China). Only clones with nonsense mutation were retained. Sequences of sgRNAs are shown in **Table S5**.

### CRISPR interference

Oligos containing sequences for sgRNAs (**Table S5**) were cloned into pLentiGuide construct (RRID:Addgene_46911)^21^. Lentiviral particles were generated by co-transfecting LentiGuide-sgRNA constructs and packaging constructs (psPAX2, RRID:Addgene_12260; and pMD2.G, RRID:Addgene_12259) to 293T cells. Supernatants with lentiviral particles were filtered through 0.45 μm mesh and added at 1:5 (vol/vol) ratio to a NB4 cell line constitutively expressing dCas9-KRAB-P2A-BFP^19^. Stably infected cells were selected with 0.5 μg/mL puromycin for one week. Enhancer knockdown efficiency and phenotype were examined by RNA-seq analysis and Giemsa stains.

### Construction of SPI1 and CEBPE overexpression plasmids

Full-length coding sequence (CDS) of human SPI1 or CEBPE was amplified from cDNA of 6th day of ATRA treated NB4. The CDS’ were then used to replace the KRAB-dCas9-DHFR coding region in an inducible-expression lentiviral plasmid (RRID:Addgene_167935) by Gibson assembly (Vazyme, Cat#C115). For the construction of SPI1 and CEBPE co-expression plasmid, P2A cleavage sequence was inserted to coding sequence by PCR primer-overlapping extension.

### RT-qPCR

Cells were harvested and resuspended immediately in RNAprotect Cell Reagent (Qiagen, Cat# 76526) for RNA stabilization. Total RNA was extracted using RNeasy Mini Kit (Qiagen, Cat# 74106) according to the manufacturer’s instructions. For each reaction, 1 μg of RNA was reverse transcribed into cDNA using HiScript III RT SuperMix (Vazyme, Cat# R323). The cDNA reaction mix was then diluted (1:10), 2 μL of which was used for one 10 μL qPCR reaction with ChamQ Universal SYBR qPCR Master Mix (Vazyme, Q711) on the LightCycler 480 II, 384 system (Roche). The relative expression level of each gene was calculated by the 2^-△△Ct^ method with GAPDH as the reference gene. Sequences of PCR primers are in **Table S5.**

### Wright-Giemsa stain

Cells were processed using cytospin via centrifuge at 800 g for 5 minutes and let dry in air. The slides were stained with Wright-Giemsa stain solution kit (Baso Cat# BA-4017). Solution A was directly added to the glass slides for 1 minute followed by double volume solution B for 8 minutes. After a brief washing with tap water, the slides were dried in air for 20 minutes before microscopic examination.

## Notes

### Competing Interest Statement

The authors have declared no competing interest.

## References

1. Tamiro F, Weng AP, Giambra V. Targeting Leukemia-Initiating Cells in Acute Lymphoblastic Leukemia. Cancer Research. 2021;81(16):4165–73. 10.1158/0008-5472.Can-20-2571.

2. de The H. Differentiation therapy revisited. Nat Rev Cancer. 2018;18(2):117–27. 10.1038/nrc.2017.103.

3. McClellan JS, Dove C, Gentles AJ, Ryan CE, Majeti R. Reprogramming of primary human Philadelphia chromosome-positive B cell acute lymphoblastic leukemia cells into nonleukemic macrophages. Proceedings of the National Academy of Sciences. 2015;112(13):4074–9. 10.1158/2159-8290.CD-21-0502.

4. Linde MH, Fan AC, Köhnke T, Trotman-Grant AC, Gurev SF, Phan P, et al. Reprogramming Cancer into Antigen-Presenting Cells as a Novel Immunotherapy. Cancer Discovery. 2023;13(5):1164–85. 10.1158/2159-8290.Cd-21-0502.

5. Zimmermannova O, Ferreira AG, Pereira C-F. Orchestrating an immune response to cancer with cellular reprogramming. Genes & Immunity. 2023. 10.1038/s41435-023-00237-4.

6. Tenen DG. Disruption of differentiation in human cancer: AML shows the way. Nat Rev Cancer. 2003;3(2):89–101. 10.1038/nrc989.

7. Thomas D, Majeti R. Biology and relevance of human acute myeloid leukemia stem cells. Blood. 2017;129(12):1577–85. 10.1182/blood-2016-10-696054.

8. Daniel MG, Rapp K, Schaniel C, Moore KA. Induction of developmental hematopoiesis mediated by transcription factors and the hematopoietic microenvironment. Annals of the New York Academy of Sciences. 2019;1466(1):59–72. 10.1111/nyas.14246.

9. Rosa FF, Pires CF, Kurochkin I, Ferreira AG, Gomes AM, Palma LG, et al. Direct reprogramming of fibroblasts into antigen-presenting dendritic cells. Science Immunology. 2018;3(30). 10.1126/sciimmunol.aau4292.

10. de The H, Chomienne C, Lanotte M, Degos L, Dejean A. The t(15;17) translocation of acute promyelocytic leukaemia fuses the retinoic acid receptor alpha gene to a novel transcribed locus. Nature. 1990;347(6293):558–61. 10.1038/347558a0.

11. Warrell RP, Jr., de The H, Wang ZY, Degos L. Acute promyelocytic leukemia. N Engl J Med. 1993;329(3):177–89. 10.1056/NEJM199307153290307.

12. Lo-Coco F, Avvisati G, Vignetti M, Thiede C, Orlando SM, Iacobelli S, et al. Retinoic acid and arsenic trioxide for acute promyelocytic leukemia. N Engl J Med. 2013;369(2):111–21. 10.1056/NEJMoa1300874.

13. Huang ME, et al. All-trans retinoic acid with or without low dose cytosine arabinoside in acute promyelocytic leukemia. Report of 6 cases. Chinese medical journal. 1987;100:949–53.

14. Tang Y, Tian X, Xu Z, Cai J, Liu H, Liu N, et al. Induced lineage promiscuity undermines the efficiency of all-trans-retinoid-acid-induced differentiation of acute myeloid leukemia. iScience. 2021;24(5):102410. 10.1016/j.isci.2021.102410.

15. Lareau CA, Duarte FM, Chew JG, Kartha VK, Burkett ZD, Kohlway AS, et al. Droplet-based combinatorial indexing for massive-scale single-cell chromatin accessibility. Nat Biotechnol. 2019;37(8):916–24. 10.1038/s41587-019-0147-6.

16. Granja JM, Corces MR, Pierce SE, Bagdatli ST, Choudhry H, Chang HY, et al. ArchR is a scalable software package for integrative single-cell chromatin accessibility analysis. Nat Genet. 2021;53(3):403–11. 10.1038/s41588-021-00790-6.

17. Pliner HA, Packer JS, McFaline-Figueroa JL, Cusanovich DA, Daza RM, Aghamirzaie D, et al. Cicero Predicts cis-Regulatory DNA Interactions from Single-Cell Chromatin Accessibility Data. Mol Cell. 2018;71(5):858–71 e8. 10.1016/j.molcel.2018.06.044.

18. Liang C, Qiao G, Liu Y, Tian L, Hui N, Li J, et al. Overview of all-trans-retinoic acid (ATRA) and its analogues: Structures, activities, and mechanisms in acute promyelocytic leukaemia. European Journal of Medicinal Chemistry. 2021;220. 10.1016/j.ejmech.2021.113451.

19. Tan Y, Wang X, Song H, Zhang Y, Zhang R, Li S, et al. A PML/RARalpha direct target atlas redefines transcriptional deregulation in acute promyelocytic leukemia. Blood. 2021;137(11):1503–16. 10.1182/blood.2020005698.

20. McLean CY, Bristor D, Hiller M, Clarke SL, Schaar BT, Lowe CB, et al. GREAT improves functional interpretation of cis-regulatory regions. Nat Biotechnol. 2010;28(5):495–501. 10.1038/nbt.1630.

21. Gilbert LA, Larson MH, Morsut L, Liu Z, Brar GA, Torres SE, et al. CRISPR-mediated modular RNA-guided regulation of transcription in eukaryotes. Cell. 2013;154(2):442–51. 10.1016/j.cell.2013.06.044.

22. Panigrahi A, O’Malley BW. Mechanisms of enhancer action: the known and the unknown. Genome Biol. 2021;22(1):108. 10.1186/s13059-021-02322-1.

23. Alon U. Network motifs: theory and experimental approaches. Nat Rev Genet. 2007;8(6):450–61. 10.1038/nrg2102.

24. McKenzie MD, Ghisi M, Oxley EP, Ngo S, Cimmino L, Esnault C, et al. Interconversion between Tumorigenic and Differentiated States in Acute Myeloid Leukemia. Cell Stem Cell. 2019;25(2):258–72 e9. 10.1016/j.stem.2019.07.001.

25. Steen CB, Liu CL, Alizadeh AA, Newman AM. Profiling Cell Type Abundance and Expression in Bulk Tissues with CIBERSORTx. Methods Mol Biol. 2020;2117:135–57. 10.1007/978-1-0716-0301-7_7.

26. Ye C, Ho DJ, Neri M, Yang C, Kulkarni T, Randhawa R, et al. DRUG-seq for miniaturized high-throughput transcriptome profiling in drug discovery. Nature Communications. 2018;9(1). 10.1038/s41467-018-06500-x.

27. Li J, Ho DJ, Henault M, Yang C, Neri M, Ge R, et al. DRUG-seq Provides Unbiased Biological Activity Readouts for Neuroscience Drug Discovery. ACS Chemical Biology. 2022;17(6):1401–14. 10.1021/acschembio.1c00920.

28. Spitz F, Furlong EE. Transcription factors: from enhancer binding to developmental control. Nat Rev Genet. 2012;13(9):613–26. 10.1038/nrg3207.

29. Barolo S, Posakony JW. Three habits of highly effective signaling pathways: principles of transcriptional control by developmental cell signaling. Genes Dev. 2002;16(10):1167–81. 10.1101/gad.976502.

30. Hromas R, et al. Hematopoietic lineage- and stage-restricted expression of the ETS oncogene family member PU.1. Blood. 1993;82(10):2998–3004.

31. Park DJ, Chumakov AM, Vuong PT, Chih DY, Gombart AF, Miller WH, Jr., et al. CCAAT/enhancer binding protein epsilon is a potential retinoid target gene in acute promyelocytic leukemia treatment. J Clin Invest. 1999;103(10):1399–408. 10.1172/JCI2887.

32. Weirauch MT, Yang A, Albu M, Cote AG, Montenegro-Montero A, Drewe P, et al. Determination and inference of eukaryotic transcription factor sequence specificity. Cell. 2014;158(6):1431–43. 10.1016/j.cell.2014.08.009.

33. Lanotte M, et al. NB4, a maturation inducible cell line with t(15;17) marker isolated from a human acute promyelocytic leukemia (M3). Blood. 1991;77:1080–6.

34. Bray NL, Pimentel H, Melsted P, Pachter L. Near-optimal probabilistic RNA-seq quantification. Nat Biotechnol. 2016;34(5):525–7. 10.1038/nbt.3519.

35. Love MI, Huber W, Anders S. Moderated estimation of fold change and dispersion for RNA-seq data with DESeq2. Genome Biol. 2014;15(12):550. 10.1186/s13059-014-0550-8.

36. Corces MR, Trevino AE, Hamilton EG, Greenside PG, Sinnott-Armstrong NA, Vesuna S, et al. An improved ATAC-seq protocol reduces background and enables interrogation of frozen tissues. Nature Methods. 2017;14(10):959–62. 10.1038/nmeth.4396.

37. Amemiya HM, Kundaje A, Boyle AP. The ENCODE Blacklist: Identification of Problematic Regions of the Genome. Scientific Reports. 2019;9(1). 10.1038/s41598-019-45839-z.

38. Schep AN, Wu B, Buenrostro JD, Greenleaf WJ. chromVAR: inferring transcription-factor-associated accessibility from single-cell epigenomic data. Nat Methods. 2017;14(10):975–8. 10.1038/nmeth.4401.

