## Supplemental Figure S1-S8, Supplemental Table S1-S7 for "Single-cell multiomics reveals a gene regulatory circuit promoting leukemia cell differentiation": supplementaryFigures_240222.pdf

Supplementary Figures S1-S8

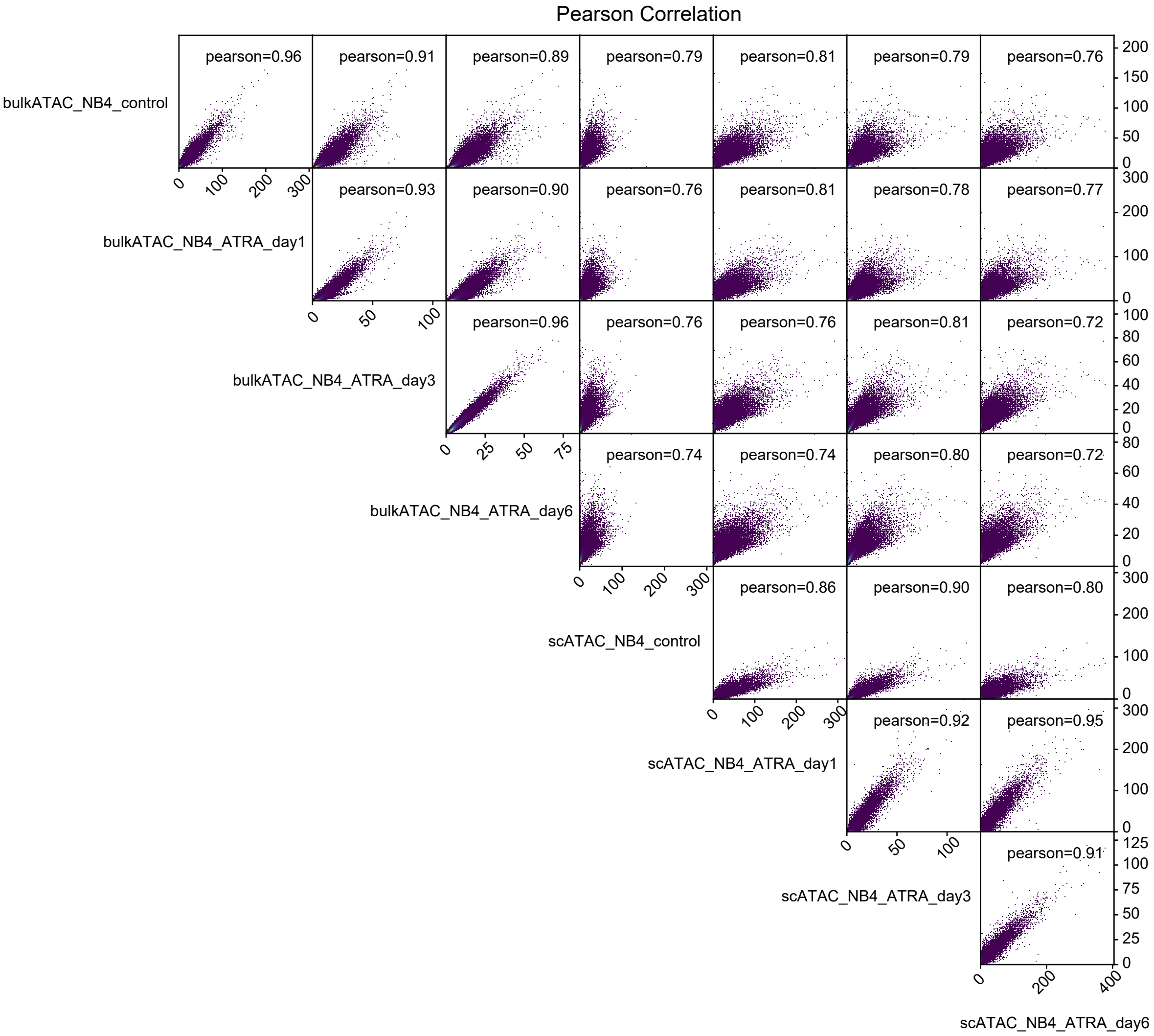


**Figure S1. Pair-wise correlations between bulk and single-cell ATAC-seq data.**

Pearson correlation coefficients are shown in each square.

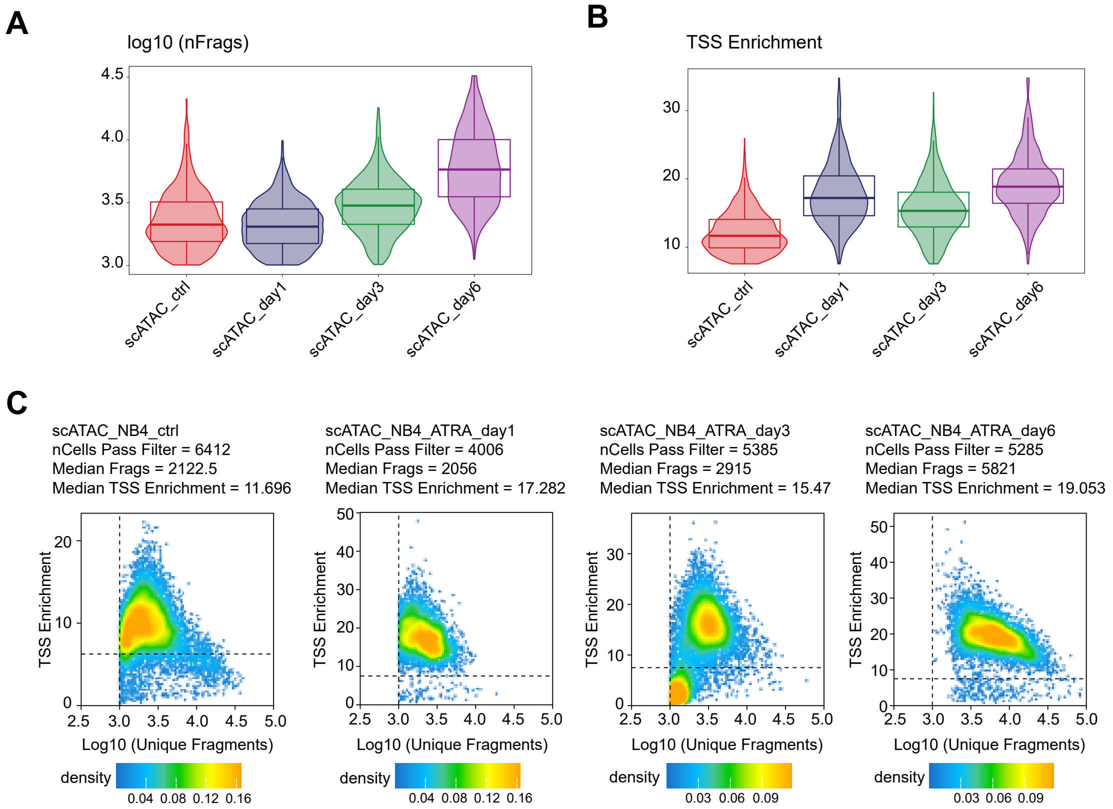


**Figure S2. Pre-processing parameters of scATAC-seq dataset.**

**(A)** Violin plots of the fragments in scATAC-seq experiments. **(B)** Violin plots of the enrichment of scATAC-seq fragments in transcription start sites (TSS). **(C)** Pseudo-color plots of filtered cells in scATAC-seq experiments.

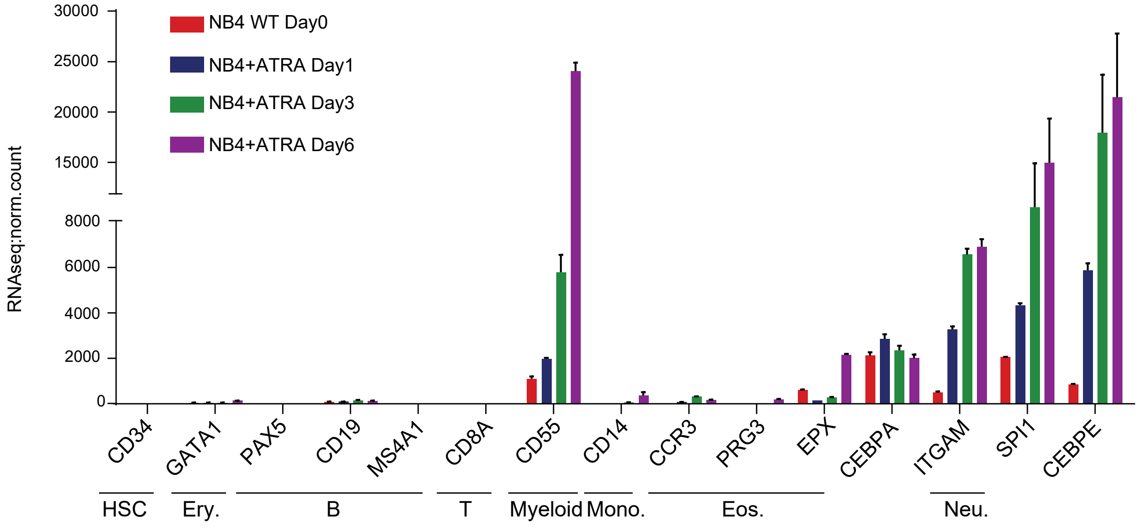


**Figure S3. Expression profiles of representative hematopoietic lineage marker genes in bulk RNA-seq experiments.**

Normalized RNA-seq counts were from DESeq2 analysis.

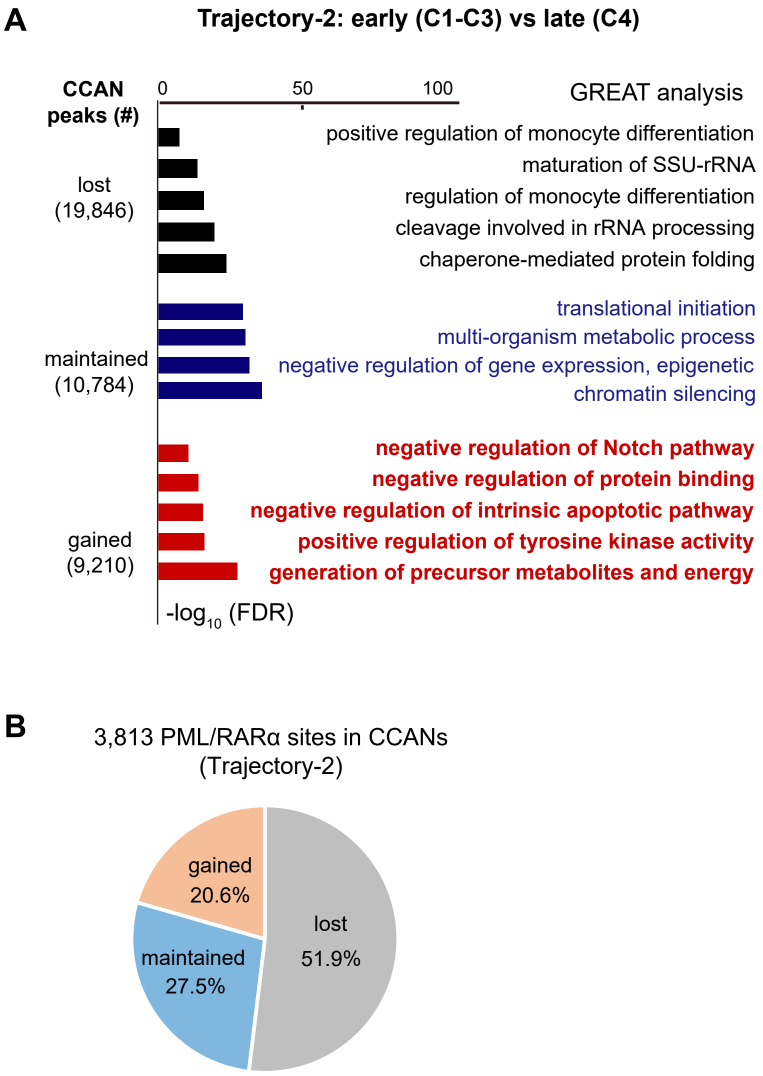


**Figure S4. Dynamics of accessible peaks along Trajectory-2.**

**(A)** Dynamics of accessible peaks during early-to-late phase transitions in Trajectory-2. The peaks that were lost, maintained , or gained during phase transitions were used as input of GREAT analysis. Bar charts of -$\log_{10}$(FDR)are shown. GO-biological process terms are at right. **(B)** The percentage of PML/RARα ChIP-seq sites in dynamic CCAN peaks during phase transitions in Trajectory-2.

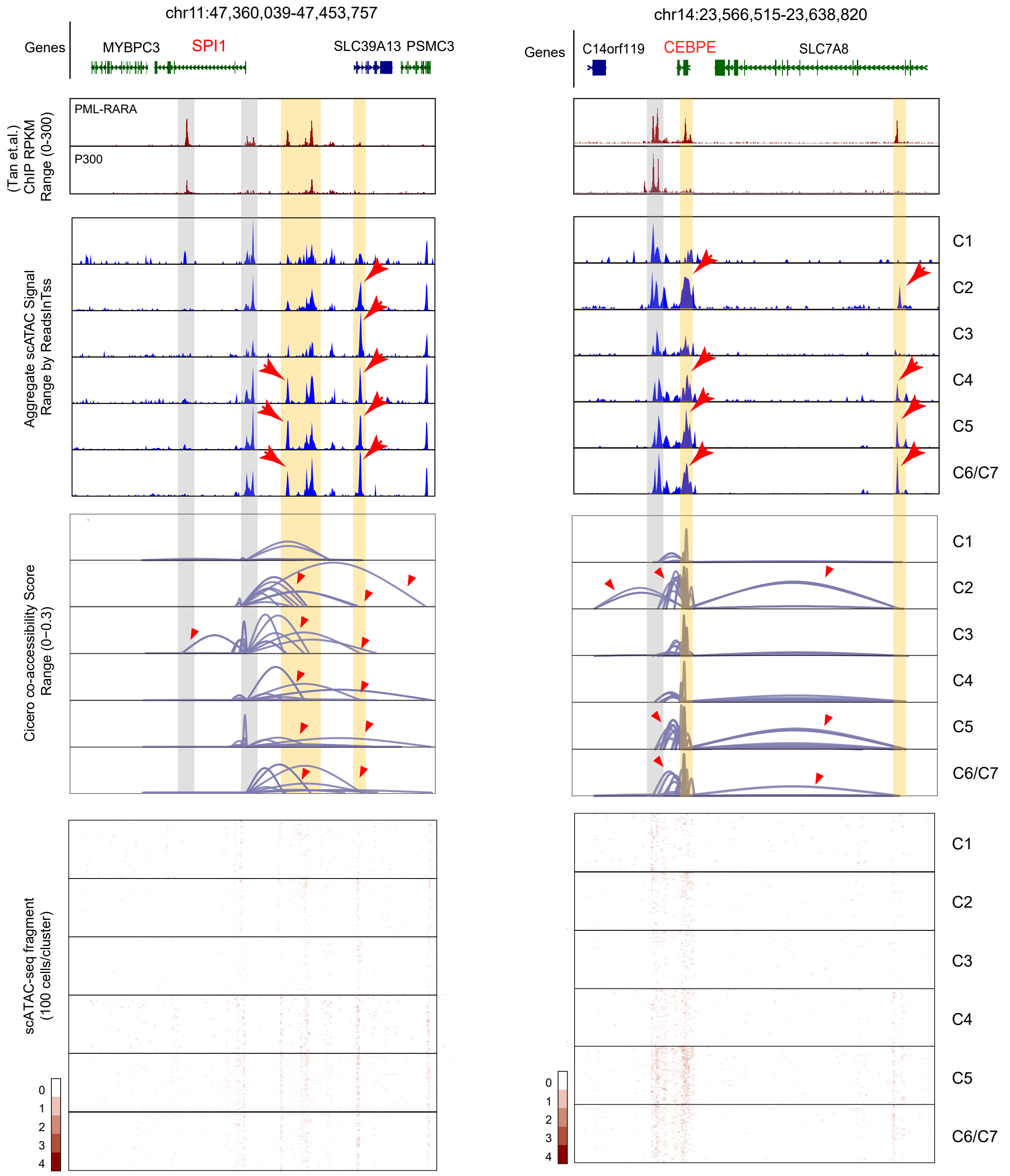


**Figure S5. Profiles of PML/RARα binding sites, scATAC-seq peaks, co-accessibility networks and frahgments at the loci of SPI1 and CEBPE.**

Notes: these tracks are the same as Figure 3G-3H, with the addition of the tracks for two clusters showing aberrant differentiaiton (C2 and C4).

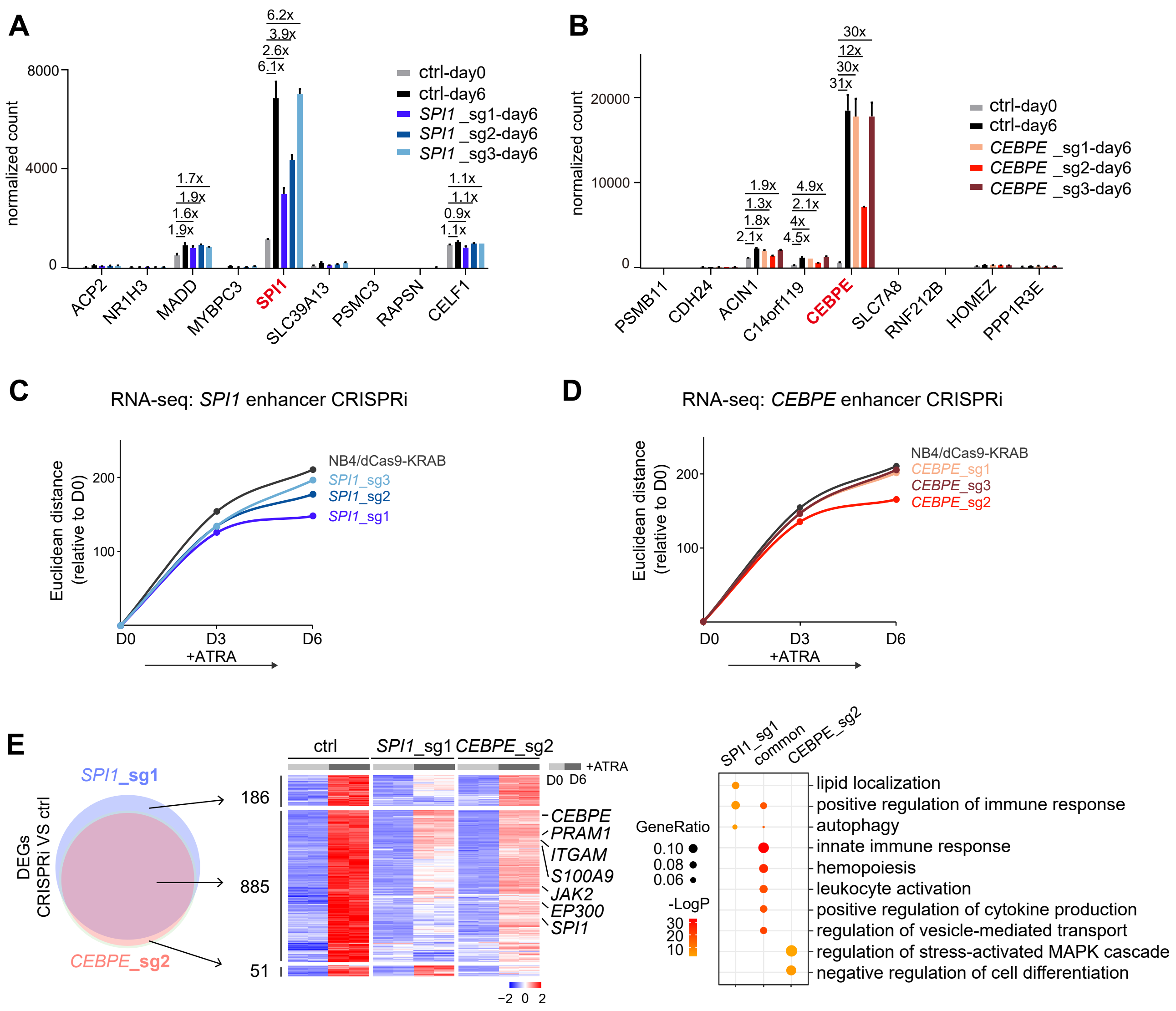


**Figure S6. RNA-seq analysis for CRISPRi experiments.**

**(A-B)** Bar plots of normlized counts of 9 genes centering on SPI1 (A) or CEBPE (B). NB4 cells expressing dCas9-KRAB and CRISPRi cells post ATRA induction are shown.

**(C-D)** Plots of Euclidean distances for control (NB4/dCas9-KRAB) and CRISPRi cell lines targeting PML/RARα binding sites near SPI1 (C) or CEBPE (D).

**(E)** Venn diagram, heatmap and GO enrichment of differentially expressed genes (DEGs) in CRISPRi experiments.

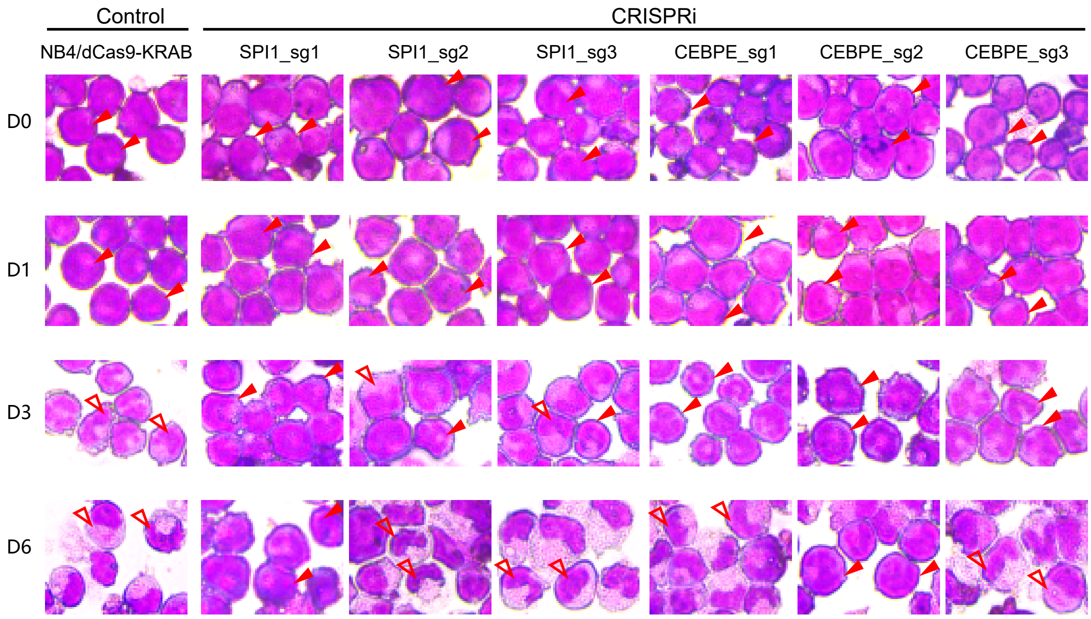


**Figure S7. Wright-Giemsa stains of CRISPRi cell lines during the course of ATRA-induction.**

Filled arrowheads indicate nuclei with blast-like characteristics. Empty arrowheads indicate condensed, multi-lobed nuclei characteristic of mature granulocytes. Horizontal axis shows different cell lines. sg: single-guide RNA. Vertical axis shows the day post ATRA-induction. D: days.

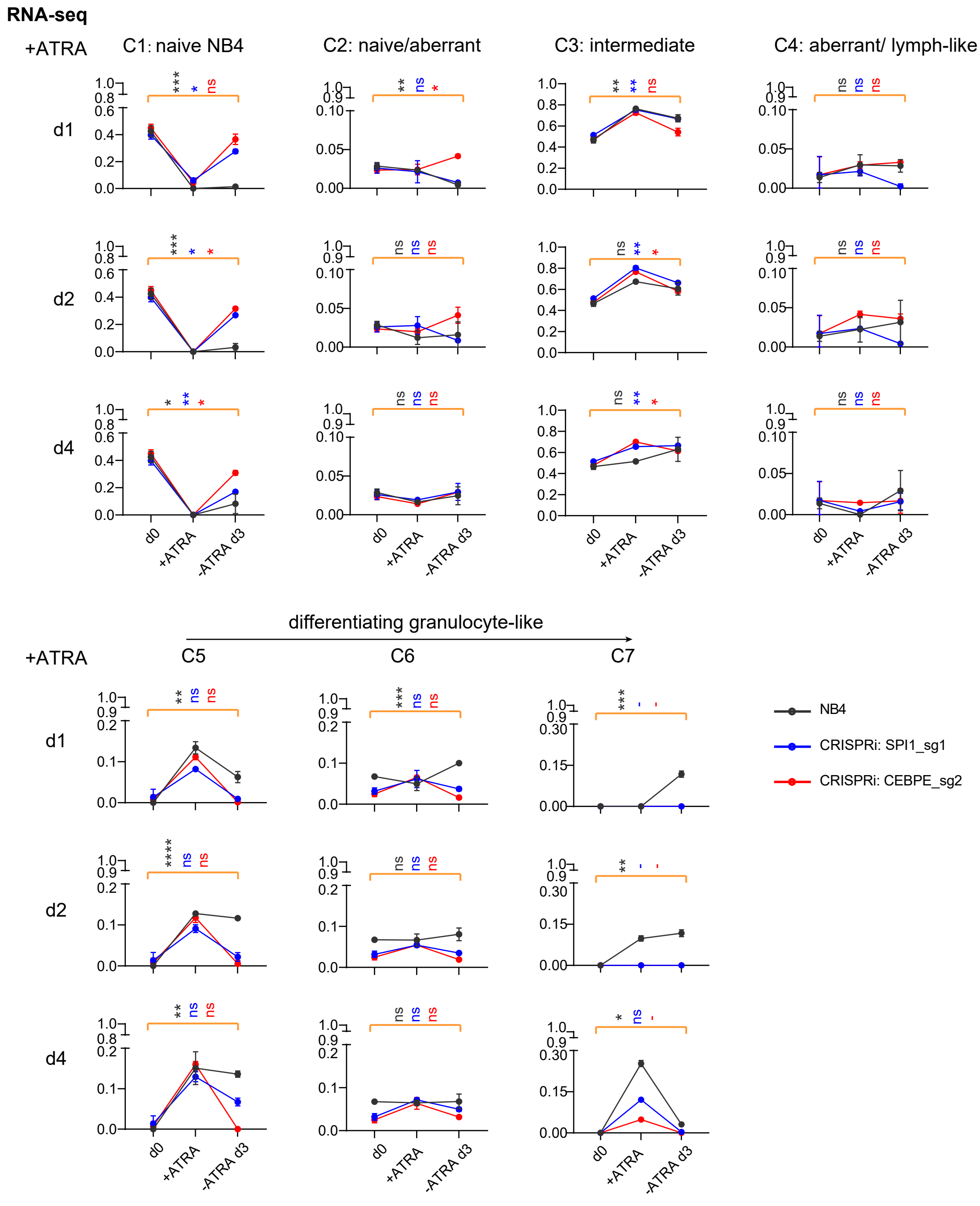


**Figure S8. CIBERSORTx analysis for bulk RNA-seq data of NB4 and CRISPRi cells in ATRA-withdrawal experiemnts.**

Cells were pre-induced with ATRA for 1/2/4 days, followed by culturing in ATRA-free medium for 3 days. The relative scores of each cluster defined by integrated scATAC-seq and scRNA-seq data are shown. Parameter t-test was conducted between each group's untreated (d0: no ATRA-induction) and ATRA-withdrawal (-ATRA d3) samples. *: $p<0.05$; **: $p<0.01$; ***: $p<0.001$; ****: $p<0.0001$; -: all values are 0.
